# Retinal *Phd2* ablation partially rescues *Crb1Crb2*-associated Leber congenital amaurosis and alters metabolic signaling

**DOI:** 10.64898/2026.10.26.761799

**Authors:** Vaishnavi Karthikeyan, Charlotte A. Andriessen, Anne Peeters, Xuefei Lu, Aat A Mulder, Zohreh Hosseinzadeh, Roman I Koning, Jan Wijnholds

## Abstract

Leber congenital amaurosis (LCA) is a severe early-onset retinal dystrophy caused by pathogenic variants in several genes, including *CRB1*. *CRB1*-associated LCA is characterized by disrupted retinal polarity, lamination, and photoreceptor organization, with no restorative treatment. We investigated whether retinal deletion of *Phd2 (Egln1)*, encoding prolyl hydroxylase domain protein 2 (PHD2), could provide mutation-independent neuroprotection in a severe developmental model of *CRB1*-associated LCA. Using conditional *Crb1/Crb2*-deficient mice, we additionally deleted *Phd2* throughout the neural retina and assessed retinal function, structure, visually guided behavior, and glycolysis-associated gene and protein expression at one and two months. Compared with *Crb1/Crb2*-deficient mice, additional *Phd2* deletion partially improved scotopic responses and significantly increased the optomotor spatial-frequency threshold, whereas b-wave and cone-mediated responses remained impaired. *Phd2Crb1Crb2*-deficient retinas displayed more organized photoreceptor rosettes containing rod- and cone-marker-positive cells, in contrast to the diffuse outer-retinal disorganization and nuclear-layer amalgamation observed in *Crb1*/*Crb2*-deficient retinas. Polarity- and synapse-associated proteins showed improved local compartmentalization within these structures. Metabolic responses were temporally selective: *Glut1* and *Vegfa* transcript levels increased at one month, whereas GLUT1 and LDHA protein levels increased at two months; *Pdk1* transcript levels decreased at 2M, whereas PDK1 protein abundance remained unchanged. Although *Phd2* deletion did not correct the underlying Crumbs-complex defect or normalize retinal lamination, and elevated *Vegfa* coincided with persistent vascular abnormalities, these findings demonstrate that retinal *Phd2* deletion partially modifies the structural and functional consequences of severe *Crb1/Crb2* deficiency and supports further investigation of PHD2-associated metabolic signaling as a potential disease-modifying target for severe *CRB1*-associated retinal degeneration.

## Introduction

Leber congenital amaurosis (LCA) is a rare group of inherited retinal dystrophies causing severe visual impairment from birth or early infancy, affecting about 1 in 40,000 newborns. It manifests with nystagmus, poor pupillary light responses, photophobia, high hyperopia, and severely reduced visual acuity and is linked to mutations in roughly two dozen genes such as *CEP290, GUCY2D, CRB1, and RPE65,* which disrupt retinal development and photoreceptor function (1–3). Pathological variations in the human *CRB1* gene account for approximately 10% of LCA cases (4). The *CRB1* gene encodes a transmembrane protein that localizes to the subapical region next to adherens junctions in mouse retinal progenitors, photoreceptors, and Müller glial cells, where it establishes and maintains apicobasal polarity for proper cell adhesion and retinogenesis (5). Through its PDZ-binding domain, CRB1 anchors the Crumbs protein complex by linking to PALS1, which recruits PATJ or MUPP1 to modulate critical pathways like Notch, Hippo, and mTORC1 that govern progenitor cell division and differentiation (6, 7).

In *CRB1*-associated disease, mouse and patient data show that loss of Crumbs-mediated polarity and adhesion disrupts lamination and photoreceptor organization during development, directly linking the gene to a LCA-like phenotype (4, 8, 9). Homozygous *Crb1* knockout mice (*Crb1*^-/-^) exhibit only mild retinal disorganization restricted to the inferior temporal quadrant (5, 10); homozygous *Crb1*^-/-^ mice with an additional single conditional retinal knockout *Crb2* allele (*Crb1*^-/-^*Crb2*^F/+^Chx10*Cre*^Tg/+^) show severe retinal disorganization restricted to the entire inferior quadrant (11); heterozygous *Crb1*^-/+^ mice with both *Crb2* alleles ablated in the retina (*Crb1*^-/+^*Crb2*^F/F^Chx10*Cre*^Tg/+^) show a phenotype throughout the retina mimicking a mild form of LCA (11); whereas combined *Crb1*/*Crb2* double homozygous conditional retinal knockout mice (*Crb1*^-/-^*Crb2*^F/F^Chx10*Cre*^Tg/+^) display profound developmental defects throughout the retina that closely recapitulate severe human *CRB1*-LCA pathology, highlighting substantial functional redundancy between the two CRB proteins (9, 12). In double-homozygous *Crb1Crb2* conditional knockout retinas, excessive progenitor proliferation, disrupted laminar organization, and a markedly thickened retinal structure mirror the severe histological and functional deficits seen in *CRB1*-linked LCA (11, 12).

Photoreceptors are among the most metabolically demanding cells in the body, consuming vast amounts of glucose to sustain phototransduction, the visual cycle, and constant membrane turnover despite their post-mitotic state (13, 14). Rods predominantly employ aerobic glycolysis (the Warburg effect), converting glucose to lactate even under oxygen-rich conditions and exporting it via the monocarboxylate transporter 1 (MCT1) to fuel Müller glia and cones, which favor oxidative phosphorylation (15). This rod-to-cone lactate shuttle maintains retinal homeostasis, with photoreceptors fully oxidizing only ∼1–3% of glucose while prioritizing rapid cytoplasmic ATP generation for outer segment renewal (16). Photoreceptors and the retinal pigment epithelium (RPE) engage in coordinated metabolic interplay: the RPE transports glucose from the choroidal circulation to photoreceptors while preferentially using photoreceptor-derived lactate and fatty acids as mitochondrial substrates, preserving glucose for photoreceptor glycolysis (17). This reciprocal flux helps coordinate redox balance and mitochondrial activity across the outer retina, limiting oxidative stress and nutrient competition (18, 19). Cell polarity defects compromise the rod-cone lactate shuttle essential for retinal metabolism, inducing glycolytic collapse, mitochondrial overload, and rapid degeneration despite preserved outer limiting membrane fragments (20, 21).

Metabolic reprogramming has emerged as a promising strategy to restore retinal structure and function in degenerative and ischemic conditions aiming to rebalance cellular energy metabolism and redox homeostasis to support neuronal survival across genetically distinct models of degeneration (22). Modulation of the PHD2/HIF-1α protein axis exemplifies the therapeutic potential of metabolic reprogramming in retinal degeneration. In the *Pde6b* mouse model of autosomal-recessive retinitis pigmentosa, rod-specific genetic ablation of *Phd2* (*Egln1*) stabilizes HIF signaling and induces a glycolytic transcriptional program, including upregulation of *Ldha* and *Pdk1*, preserving anabolic flux and leading to robust rod preservation, secondary cone survival, and sustained visual function. Extending these findings, AAV-mediated CRISPR editing of *Phd2* in rods similarly rescued photoreceptors in both recessive and dominant retinitis pigmentosa (RP) models, showing that reprogramming aerobic glycolysis is a gene-agnostic strategy for overcoming genetic heterogeneity in retinal degeneration (23). Likewise, SIRT6 inhibition reprograms rod metabolism toward increased glycolytic flux, promoting rod and cone survival in *PDE6*-associated degeneration models (24). Collectively, these studies support metabolic rewiring at the glycolysis–mitochondrial interface as a mutation-independent strategy for sustaining photoreceptor viability.

To investigate whether *Phd2*-mediated metabolic reprogramming confers broader neuroprotection *in vivo*, we extended this approach to a previously characterized genetic mouse model of *Crb1*-associated LCA (12). Unlike the later-onset, rod-centric degeneration of the *Pde6β*- and *Rho*-based RP models used in prior studies, this model exhibits early-onset developmental defects, aberrant lamination, and expanded progenitor-cell proliferation that closely recapitulate the severe pathology of human *CRB1*-LCA. The disease model carries a homozygous *Crb1* knockout and a conditional deletion of *Crb2* exons 10–13, which are flanked by *loxP* sites. In retinal progenitor cells, Cre recombinase is expressed under control of the Chx10 promoter, resulting in *Crb2* excision (*Crb1*^−/−^*Crb2*^F/F^Chx10*Cre*^Tg/+^; here abbreviated as *Crb1Crb2-*cKO) (25); given the redundancy between Crb1 and Crb2 proteins, loss of Crb2 on a *Crb1*-null background is required to reveal the full phenotype severity. Control littermates lacking the *Cre* driver (*Crb1Crb2*) retained intact *Crb2* alleles and displayed only the mild inferior-quadrant disorganization characteristic of *Crb1* single mutants, serving as the phenotypic baseline on a C57BL/6J background. To assess whether pan-retinal HIF stabilization can ameliorate this severe developmental degeneration, we additionally deleted *Egln1* (*Phd2*) throughout the neural retina using the same Chx10-*Cre* driver, yielding a putative rescue genotype (*Phd2*^F/F^*Crb1*^−/−^*Crb2*^F/F^Chx10*Cre*^Tg/+^ (12, 26); here abbreviated as *Phd2Crb1Crb2-*cKO). In parallel, we investigated whether PHD2 protein loss remodels retinal glycolytic metabolism and whether these changes contribute to structural and functional rescue. Collectively, this study evaluates metabolic reprogramming as a mutation-independent therapeutic strategy in a physiologically relevant model of severe *CRB1*-LCA.

## Results

### Phd2 deletion improves photoreceptor-driven retinal function in severe Crb1Crb2-cKO-LCA model

To generate the three experimental genotypes as littermates, we crossed *Phd2*^F/+^*Crb1*^−/−^*Crb2*^F/F^Chx10*Cre*^Tg/+^ mice with *Phd2*^F/+^*Crb1*^−/−^*Crb2*^F/F^ mice; all lines were maintained on a C57BL/6J genetic background. The Chx10-*Cre* transgene drives Cre recombinase expression throughout the developing neural retina from embryonic day (E)11.5, in all retinal cell types except the retinal pigment epithelium (27) as previously described (28). This cross yielded, within the same litters: (i) phenotypic controls (*Crb1Crb2*, lacking Chx10-*Cre*; here abbreviated as *Crb1Crb2-*control), in which *Crb2* remains intact and only the mild *Crb1*-null phenotype is manifest; (ii) phenotypic (severe LCA) mice (*Crb1*^−/−^*Crb2*^F/F^Chx10*Cre*^Tg/+^, *Phd2*^+/+^; here abbreviated as *Crb1Crb2-*cKO), in which conditional deletion of *Crb2* on the *Crb1*-null background abolishes *Crb1*/*Crb2* redundancy and reproduces the severe early-onset LCA phenotype previously characterized (11); and (iii) *Phd2*-knockout mice (*Phd2*^F/F^*Crb1*^−/−^*Crb2*^F/F^Chx10*Cre*^Tg/+^; here abbreviated as *Phd2Crb1Crb2-*cKO), in which both *Crb2* and *Phd2* are concurrently deleted throughout the neural retina. We performed *in vivo* functional and structural analysis at 1 and 2 months of age using electroretinography (ERG), spectral-domain optical coherence tomography (SD-OCT), and confocal scanning laser ophthalmoscopy (cSLO); all experiments and data analysis were performed by investigators blinded to genotype, with animals randomized and co-housed with littermates of all genotypes.

To determine whether *Phd2* deletion improves retinal function in the severe *CRB1*-LCA model, we performed scotopic and photopic ERG in *Crb1Crb2-*control, phenotypic *Crb1Crb2-*cKO, and *Phd2Crb1Crb2-*cKO mice at one month of age (29) (Figure 1A). Quantification of the scotopic a-wave intensity-response series showed that *Phd2Crb1Crb2-*cKO amplitudes trended above the phenotypic *Crb1Crb2-*cKO model across flash intensities (Figure 1B), Scotopic a-wave amplitudes were higher in *Phd2Crb1Crb2*-cKO than in *Crb1Crb2*-cKO mice across several flash intensities, reaching statistical significance at 1.5 log cd·s/m² : a-wave amplitude increased from ∼5 µV in the phenotypic model to ∼40–60 µV in the *Phd2Crb1Crb2*-cKO (p < 0.01; control ∼250 µV) (Figure 1D). This improvement in photoreceptor-driven signaling was also reflected in the b/a-wave amplitude ratio, which is markedly elevated when the a-wave is selectively lost relative to the b-wave. The b/a ratio was significantly lower in the *Phd2Crb1Crb2-*cKO than in the phenotypic model (p < 0.05), falling from a phenotypic median of ∼12 to control-like values of ∼1–2 in the rescue genotype (Figure 1F), consistent with the observed increase in a-wave amplitudes. Together, these findings demonstrate a significant improvement in scotopic photoreceptor-driven responses following *Phd2* deletion at 1M of age. Scotopic b-wave amplitude at 1.5 log cd·s/m² did not differ significantly between *Crb1Crb2-*cKO and *Phd2Crb1Crb2-*cKO genotypes (Figure 1E), and the full scotopic b-wave intensity-response series showed a comparable degree of impairment in both genotypes relative to control (Figure 1C). Photopic b-wave and flicker ERG responses likewise remained comparably depressed in the *Crb1Crb2-*cKO, and *Phd2Crb1Crb2-*cKO relative to control, with no distinguishable improvement conferred by *Phd2* deletion at these cone-driven readouts (Figure 1G–H).

**Figure 1:**
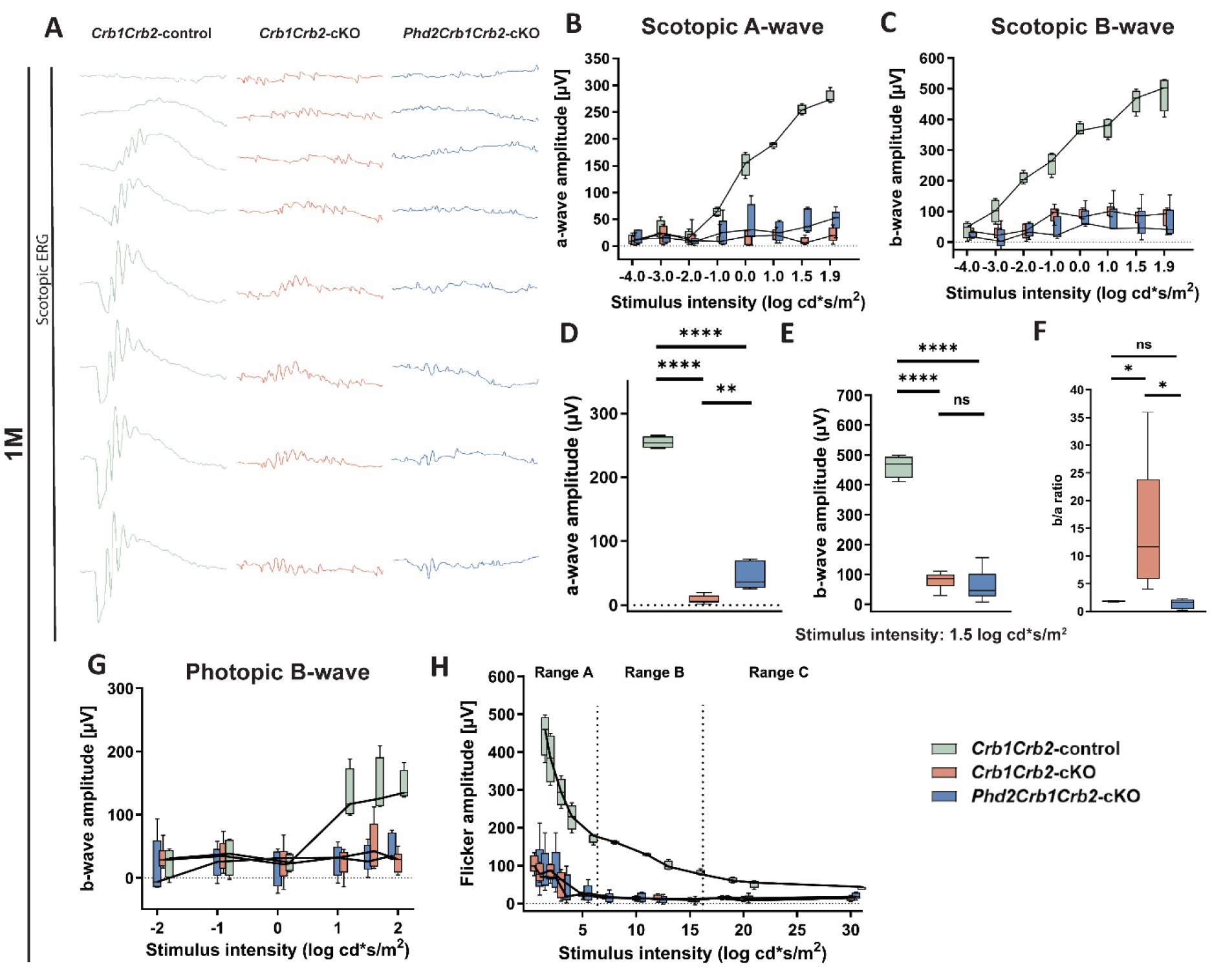
ERG analysis reveals partial improvement of photoreceptor-driven retinal function following *Phd2* deletion at one month of age. Retinal function in *Crb1Crb2*-control, *Crb1Crb2*-cKO, and *Phd2Crb1Crb2*-cKO mice was assessed by single-flash and flicker electroretinography at 1 month of age. (A) Representative scotopic single-flash ERG traces recorded from the indicated groups. (B,C) Scotopic a-wave and b-wave amplitudes plotted as a function of the logarithm of flash intensity. (D–F) Quantification of scotopic a-wave amplitude, b-wave amplitude, and b-wave/a-wave ratio at 1.5 log cd·s/m². (G) Photopic b-wave amplitudes plotted as a function of the logarithm of flash intensity. (H) Flicker ERG amplitudes plotted as a function of stimulus intensity. *Crb1Crb2*-control, n=4; *Crb1Crb2*-cKO, n=6; *Phd2Crb1Crb2*-cKO, n=5 animals. Boxes indicate the interquartile range between 25% and 75%, whiskers indicate the 5% and 95% quantiles, and central lines indicate the median. Statistical comparisons among genotypes were performed using one-way ANOVA followed by Tukey’s multiple-comparisons test. **** *p*<0.0001, ** *p*<0.001, * *p*<0.05, *ns* not significant

To investigate whether the improvement in photoreceptor-driven retinal responses following *Phd2* deletion translated into improved visual performance, optomotor responses were evaluated in 1-month-old mice. Contrast sensitivity increased with spatial frequency in all experimental groups. *Phd2Crb1Crb2-*cKO mice showed an upward shift in contrast-sensitivity responses relative to *Crb1Crb2-*cKO mice, most prominently at 0.064 and 0.092 cycles/degree (Fig. 2A). Consistent with this pattern, *Phd2* deletion significantly increased spatial-frequency threshold relative to *Crb1Crb2-*cKO mice (Fig. 2B). These data show that *Phd2* deletion improves optomotor visual function in the severe *Crb1Crb2-*cKO model at 1 month of age and corroborate the improvement in photoreceptor-driven retinal function detected by ERG, consistent with reports that human *CRB2*-directed viral gene therapy improves both ERG and optomotor-based contrast sensitivity in *Crb1*/*Crb2* mouse models (30, 31).

**Figure 2.**
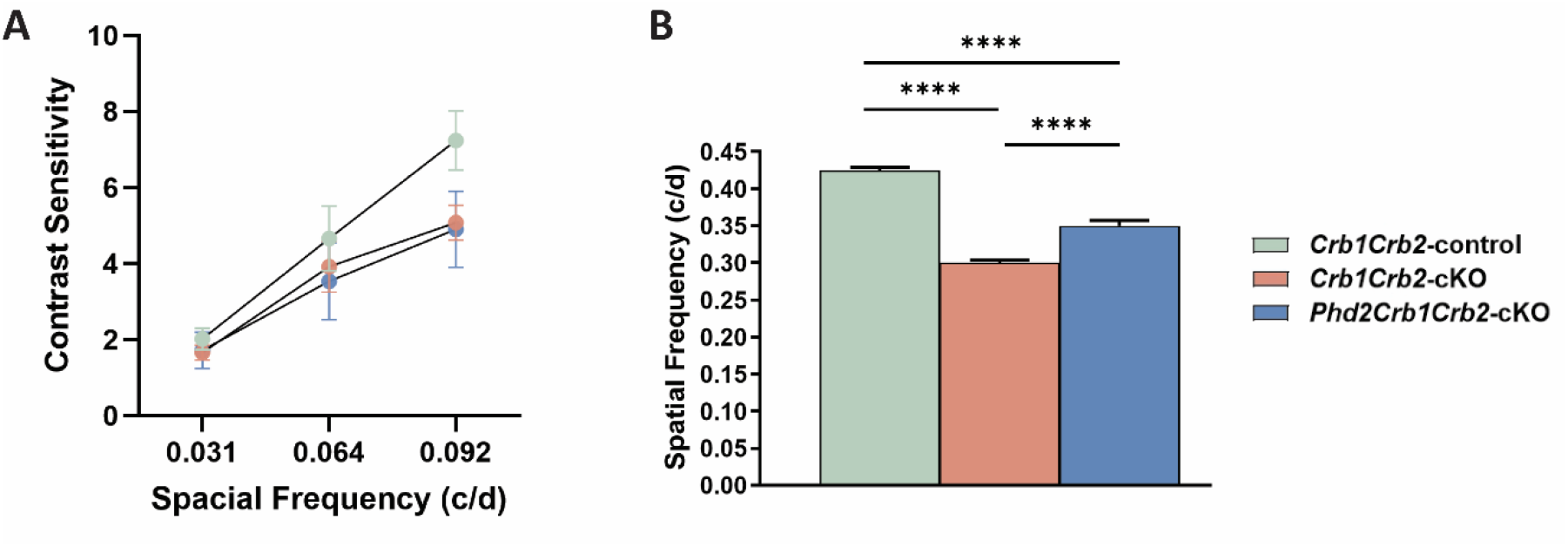
*Phd2* deletion improves optomotor visual function in the severe *Crb1Crb2-*cKO model at one month of age. Optomotor responses were assessed in 1-month-old C*rb1Crb2*-control, *Crb1Crb2*-cKO, and *Phd2Crb1Crb2*-cKO mice. (A) Contrast sensitivity measured at spatial frequencies of 0.031, 0.064, and 0.092 cycles/degree. (B) Spatial frequency threshold at 1 month of age. *Phd2* deletion increased contrast sensitivity and significantly improved the spatial frequency threshold relative to *Crb1Crb2-*cKO mice. *Crb1Crb2*-control, n=13; *Crb1Crb2*-cKO, n=14; *Phd2Crb1Crb2*-cKO, n=14. Data are presented as mean ± SEM. Statistical comparisons among genotypes were performed using one-way ANOVA followed by Tukey’s multiple-comparisons test. **** *p*<0.0001

### Phd2 deletion alters the pattern of outer retinal remodeling in the severe Crb1Crb2-cKO-LCA model

*In vivo* retinal morphology was assessed using confocal scanning laser ophthalmoscopy (cSLO) for fundus imaging, fluorescein angiography (FA) for retinal vasculature, and spectral-domain optical coherence tomography (SD-OCT) for cross-sectional analysis of retinal lamination, in 1- and 2-month-old *Crb1Crb2-*cKO and *Phd2Crb1Crb2-*cKO mice, and age-matched *Crb1Crb2-*controls. At both ages, *Crb1Crb2-*cKO mice displayed a granular, mottled fundus, attenuated and tortuous retinal vessels, and collapse of the outer retina into an undifferentiated hyperreflective band on SD-OCT (Fig. 3B, B′, B″, E, E′, E″). *Phd2Crb1Crb2*-cKO mice retained an abnormal fundus and retinal vasculature but, at both ages, showed discrete circular areas of increased fluorescence that were not evident in *Crb1Crb2*-cKO mice (Fig. 3C′, F′). SD-OCT similarly revealed circumscribed gaps with hyperreflective margins within the neurosensory retina, contrasting with the diffuse, undifferentiated outer-retinal collapse observed in *Crb1Crb2*-cKO mice (Fig. 3C″, F″). These distinct focal abnormalities prompted further histological characterization. *Crb1Crb2-*controls showed normal fundus, vasculature, and retinal lamination throughout (Fig. 3A, A′, A″, D, D′, D″).

**Figure 3.**
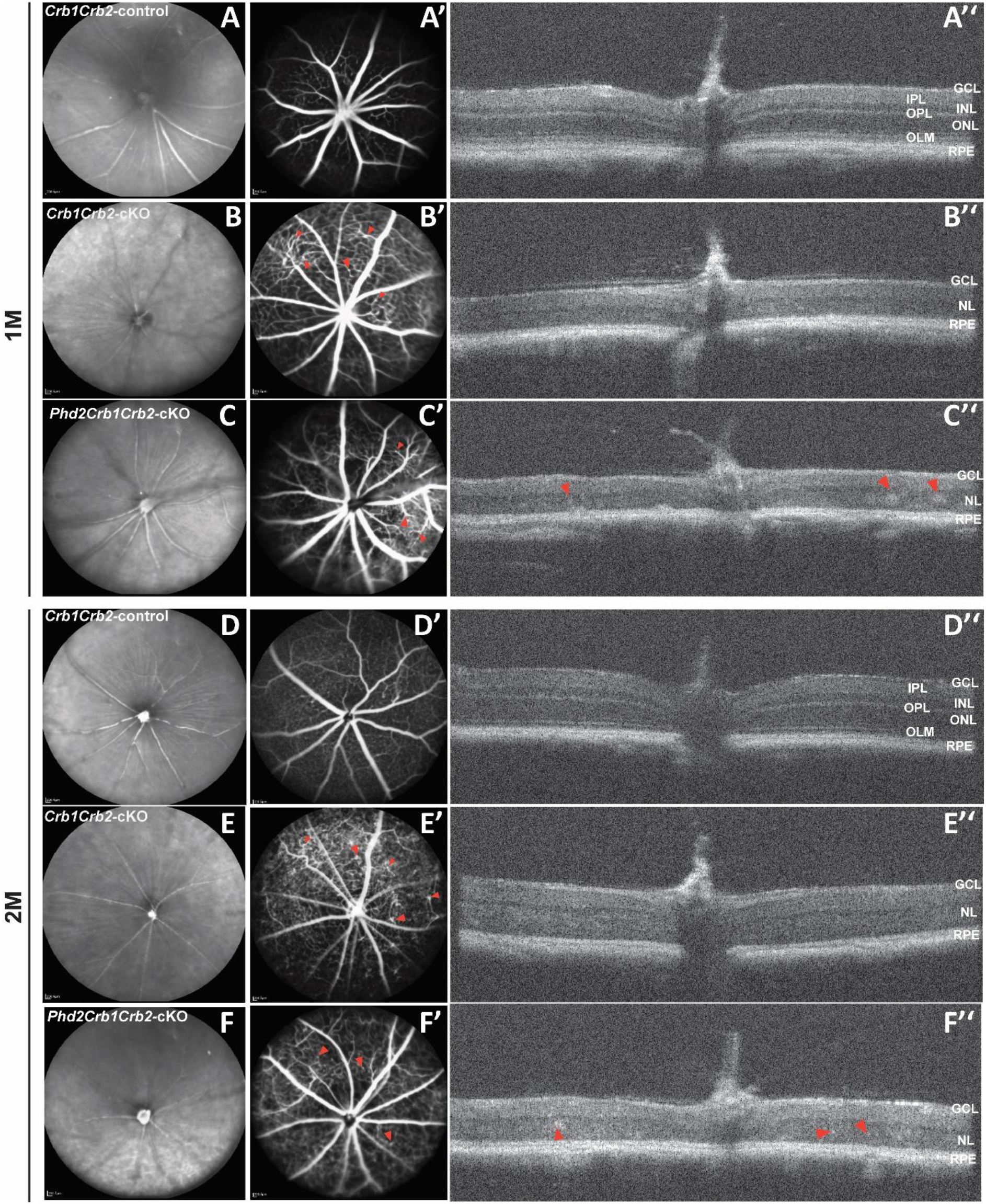
*Phd2* deletion is associated with a distinct, more contained pattern of outer retinal remodeling relative to the phenotypic severe *Crb1Crb2-*cKO model. Fundus photography (A–F), fluorescein angiography (A′–F′), and spectral-domain OCT (A″–F″) images from *Crb1Crb2-*control (A– A″, D–D″), *Crb1Crb2-*cKO (B–B″, E–E″), and *Phd2Crb1Crb2-*cKO (C–C″, F–F″) mice at 1 month (A–C″) and 2 months (D–F″) of age. Arrowheads in B′, C′, E′, and F′ indicate retinal vessel abnormalities. Arrowheads in C″ and F″ indicate discrete subretinal hyperreflective deposits characteristic of *Phd2Crb1Crb2-*cKO mice, contrasting with the diffuse outer retinal collapse seen in *Crb1Crb2-*cKO mice. All cSLO images are shown in the standard acquisition orientation, with dorsal/superior retina at the top and ventral/inferior retina at the bottom. GCL, ganglion cell layer; IPL, inner plexiform layer; INL, inner nuclear layer; OPL, outer plexiform layer; ONL, outer nuclear layer; OLM, outer limiting membrane; RPE, retinal pigment epithelium.

### Phd2 deletion promotes photoreceptor rosette organization in the severe Crb1Crb2-cKO-LCA model

To determine the cellular basis of the focal outer retinal abnormalities detected by SD-OCT, histological sections were analyzed at postnatal day 15 (P15) and 1 month (1M) of age. In *Crb1Crb2-*control retinas, the laminar architecture was preserved, with clearly distinguishable outer nuclear layer (ONL) and inner nuclear layer (INL) separated by the outer plexiform layer, and a well-defined ganglion cell layer (GCL) (Fig. 4A, A′, D, D′). In *Crb1Crb2-*cKO retinas, the outer retinal layers were severely disrupted, with the ONL and INL merging into a single, indistinct nuclear layer and consequent loss of clear laminar boundaries (Fig. 4B, B′, E, E′), consistent with the ONL–INL amalgamation described in advanced photoreceptor degenerations (32, 33). Within this disorganized tissue, rosette-like structures were present but were poorly circumscribed, indicative of an aberrant, incompletely organized clustering of photoreceptor cells.

**Figure 4.**
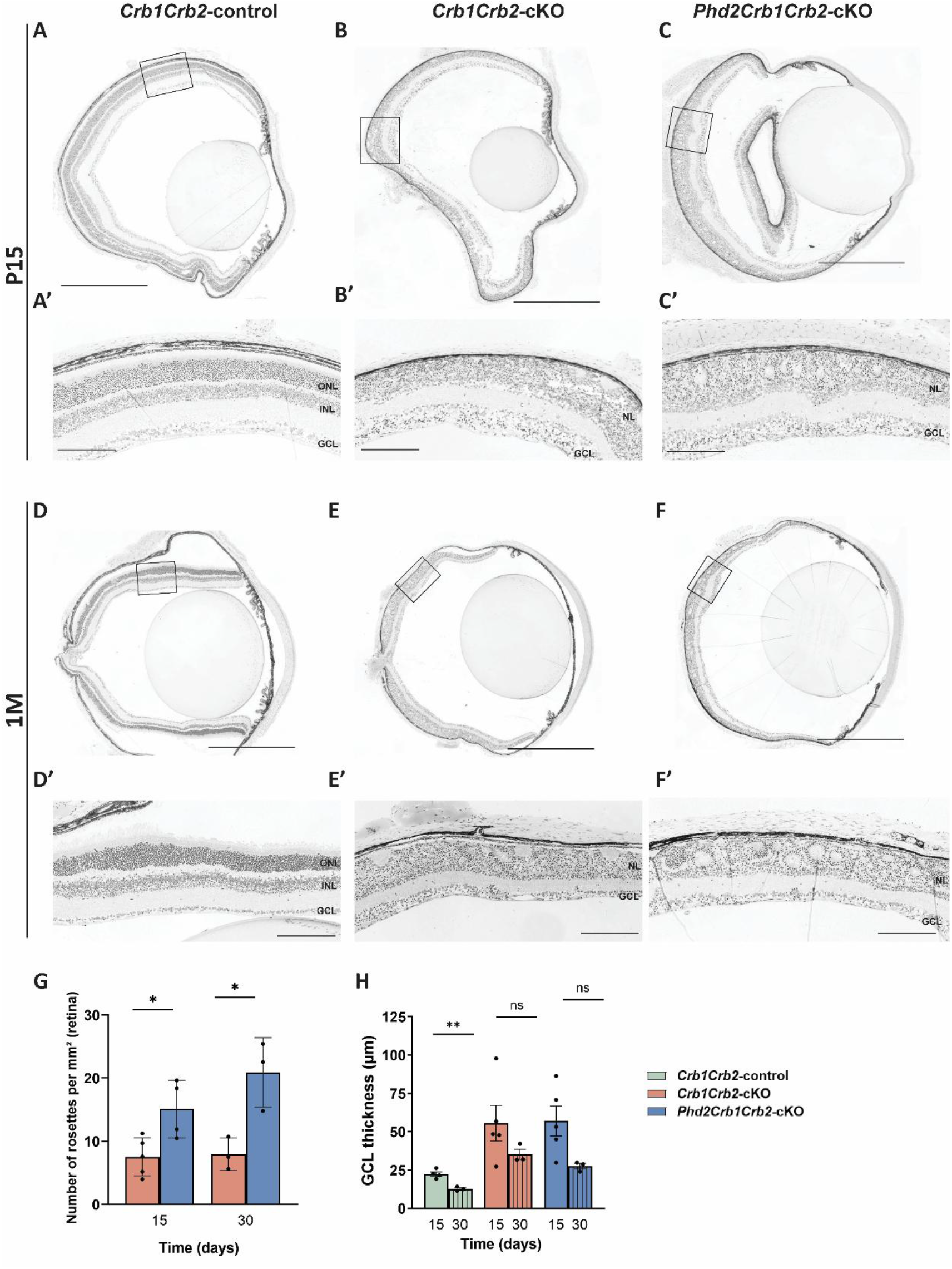
*Phd2* deletion improves photoreceptor rosette organization in the severe *Crb1Crb2-*cKO model. Histological cross-sections of *Crb1Crb2-*control (A, A′, D, D′), *Crb1Crb2-*cKO (B, B′, E, E′), and *Phd2Crb1Crb2-*cKO (C, C′, F, F′) retinas at postnatal day 15 (P15; A–C′) and 1 month (1M; D–F′) of age. Boxed regions in A–F are shown at higher magnification in A′–F′. (A–F); Scale bars: 50 µm (A′–F′). (G) Quantification of the number of ONL rosette-like structures per mm^2^ of retina in *Phd2Crb1Crb2-*cKO mice at P15 and 1M. (H) Quantification of GCL thickness (µm) in *Crb1Crb2-*control (green), *Crb1Crb2-*cKO (red), and *Phd2Crb1Crb2-*cKO (blue) mice at P15 and 1M. ONL, outer nuclear layer; INL, inner nuclear layer; GCL, ganglion cell layer. Scale bars: 500 µm. Data are presented as mean ± SEM; individual data points represent individual animals. Statistical comparisons among genotypes were performed using one-way ANOVA followed by Tukey’s multiple-comparisons test. * *p*<0.05, ** *p*<0.001, *ns* not significant

In contrast, *Phd2Crb1Crb2-*cKO retinas displayed clearly delineated separation of ONL cells into well-defined rosette structures, which were considerably more distinct than those observed in *Crb1Crb2-*cKO retinas (Fig. 4C, C′, F, F′). These rosettes corresponded histologically to the discrete hyperreflective gaps previously identified by SD-OCT, in agreement with prior SD-OCT-histology validation in *Crb1*-deficient retinas (34) (Fig. 3C″, F″). Their regular, circumscribed architecture indicates that *Phd2* deletion alters the spatial organization of photoreceptor-containing structures within the severely dysplastic retina in the severe *CRB1*-LCA model (35).

Quantification of ONL rosette-like structures showed an increase in rosette number in *Phd2Crb1Crb2-*cKO retinas compared with *Crb1Crb2-*cKO retinas at both P15 and 1 month (Fig. 4G). Ganglion cell layer thickness was also quantified, as it appeared visually reduced in *Phd2Crb1Crb2-*cKO retinas relative to *Crb1Crb2-*cKO retinas; the phenotypic group showed an apparent thickening that approached control-like values, although this difference did not reach statistical significance between *Crb1Crb2-*cKO and *Phd2Crb1Crb2-*cKO groups at either time point (Fig. 4H).

### Crb1Crb2-cKO-LCA retinas exhibit disrupted retinal lamination and photoreceptor ultrastructure

To examine the ultrastructural organization of the retina, transmission electron microscopy was performed on 1-month-old *Crb1Crb2*-control and *Crb1Crb2*-cKO retinas. *Crb1Crb2*-control retinas showed normal retinal lamination, with distinct ONL and INL organization and photoreceptor nuclei aligned in rows within the ONL (Fig. 5A). By contrast, *Crb1Crb2*-cKO retinas showed a marked loss of nuclear-layer organization, with ONL and INL nuclei intermingled within a disorganized nuclear layer. Within this layer, photoreceptor nuclei were arranged circumferentially to form rosette-like structures (Fig. 5D), consistent with the severe lamination defects previously described in this model.

**Figure 5.**
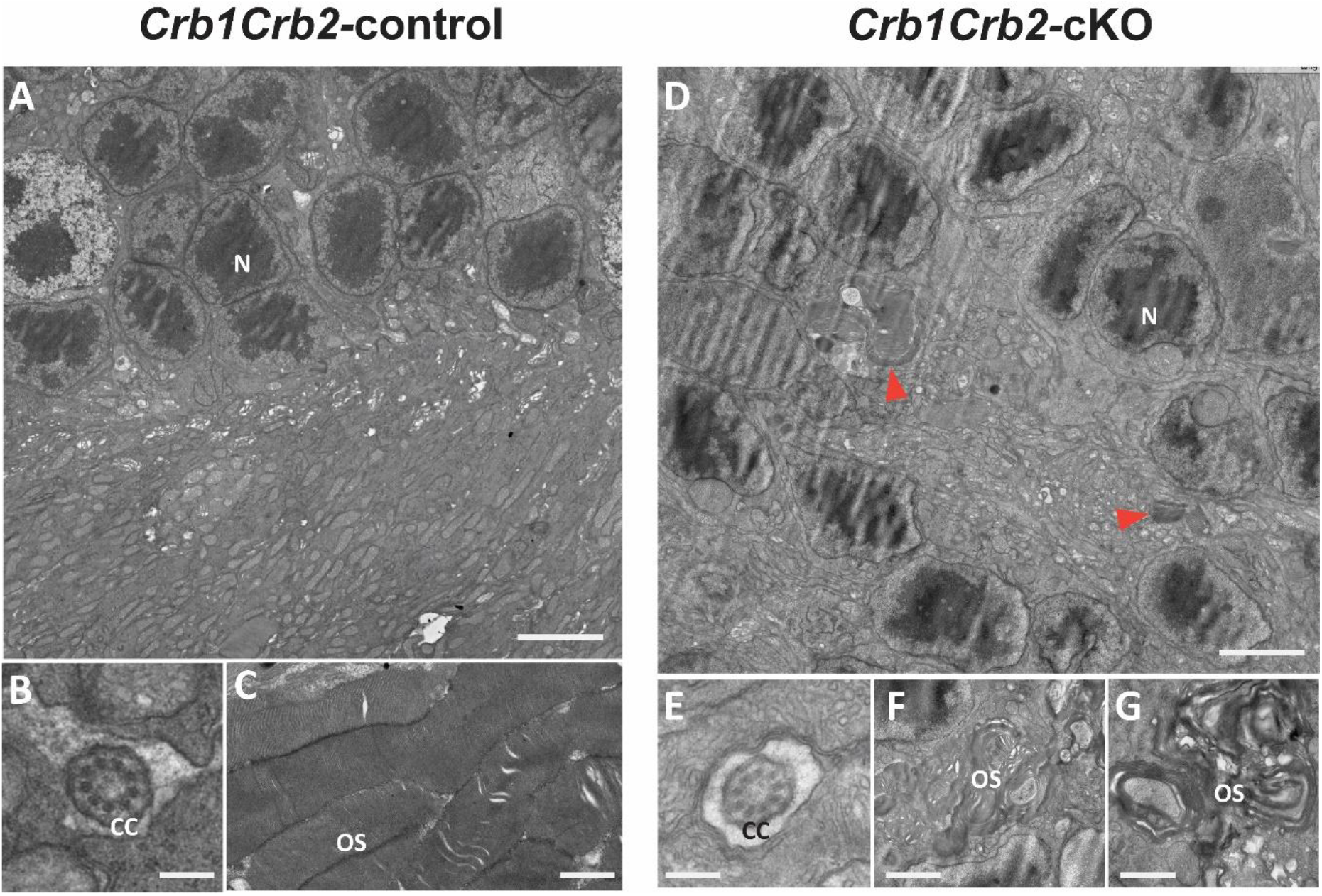
*Crb1Crb2*-cKO retinas exhibit disrupted retinal organization and photoreceptor ultrastructure. Representative transmission electron microscopy images of retinas from 1-month-old *Crb1Crb2*-control and *Crb1Crb2*-cKO mice (n = 2 eyes per genotype). In the *Crb1Crb2*-control retina, the ONL and INL were distinctly organized, with photoreceptor nuclei regularly aligned in rows within the ONL (A). Connecting cilia displayed normal morphology and localization (B), while the elongated photoreceptor outer segments contained well-organized, regularly aligned membranous discs (C). In contrast, the *Crb1Crb2*-cKO retina lacked distinct ONL and INL organization, with intermingled nuclei forming a disorganized nuclear layer and photoreceptor nuclei arranged into rosette-like structures (D). Outer segments were located within the lumen of the rosettes (D; arrowheads), where their membranous discs appeared irregularly oriented and markedly disorganized compared with those in the control retina (F, G). Connecting cilia retained an apparently normal morphology but were located within the rosette lumen (E). CC, connecting cilium; INL, inner nuclear layer; N, nucleus; ONL, outer nuclear layer; OS, outer segment. Scale bars: 3 µm (A, D); 250 nm (B, E); 1 µm (C, G); 2 µm (F).

At higher magnification, connecting cilia exhibited a normal ultrastructural appearance in both *Crb1Crb2*-control and *Crb1Crb2*-cKO retinas (Fig. 5B, E). However, whereas connecting cilia were appropriately positioned in the control retina, they were located within the centres of the rosettes in the *Crb1Crb2*-cKO retina. Control photoreceptor outer segments were elongated and contained well-organized membranous discs (Fig. 5C). In contrast, outer segments were ectopically located within the rosette centres in *Crb1Crb2*-cKO retinas and displayed irregularly oriented and disorganized disc membranes (Fig. 5D, F, G).

### Rod and cone photoreceptors are organized within rosette-like structures following Phd2 deletion

To define the cellular composition of the rosette-like structures in the outer nuclear layer (ONL), we performed immunofluorescence staining for rod and cone photoreceptor markers at postnatal day 15 (P15) and 1 month (1M) (Fig. 6a). In *Crb1Crb2*-control retinas, rhodopsin immunoreactivity was confined to rod outer and inner segments at the apical retinal surface, with DAPI-labeled nuclei forming a compact ONL above a clearly separated inner nuclear layer (INL) and ganglion cell layer (GCL), consistent with normal retinal lamination (Fig. 6a A,D). Recoverin labeled photoreceptor somata within the ONL and cone bipolar cells in the INL (Fig. 6a M,P), and red/green opsin signal was restricted to cone outer segments (Fig. 6a G,J), further confirming intact photoreceptor identity and organization.

**Figure 6a.**
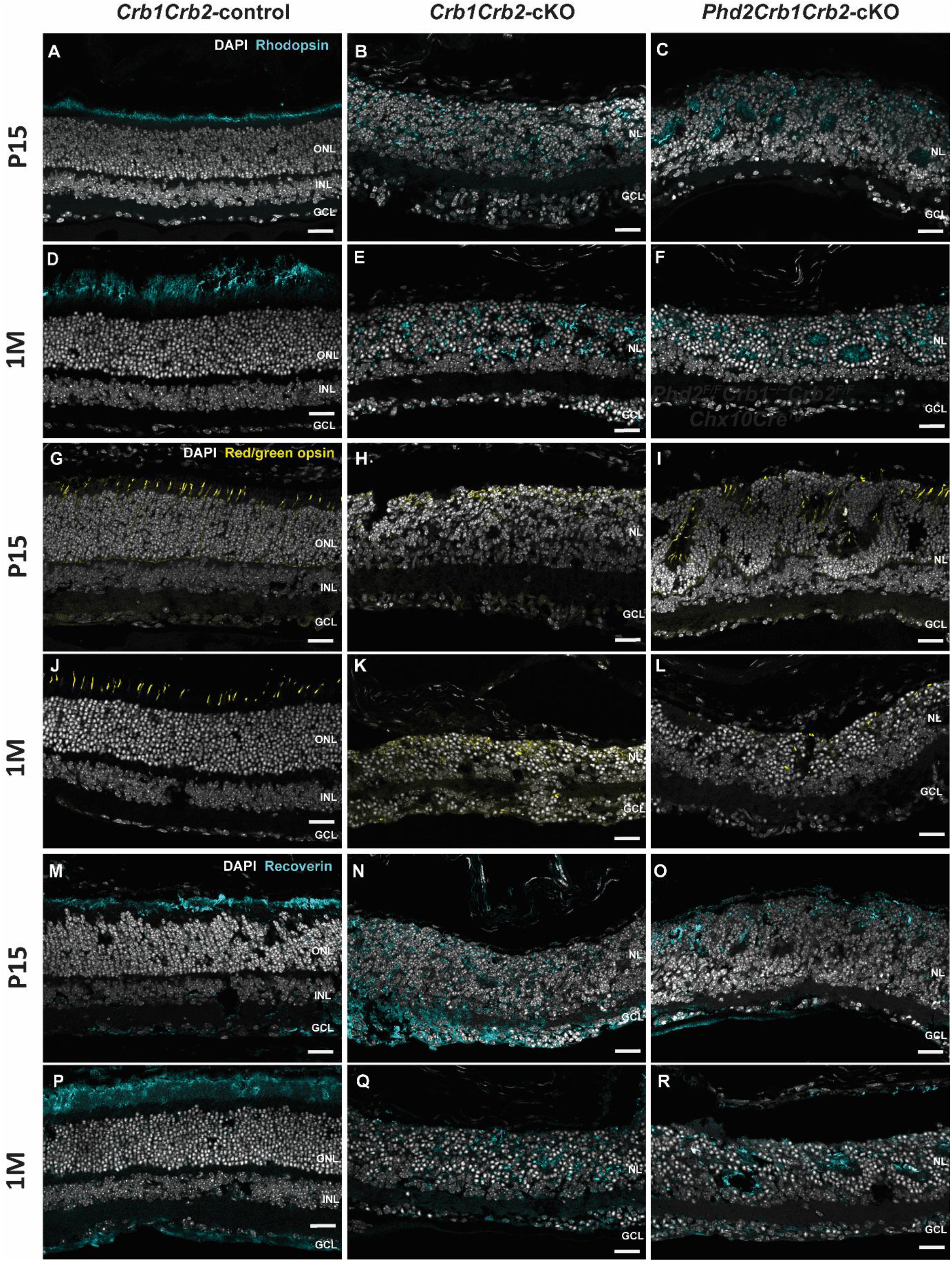
Rod and cone photoreceptors are spatially organized within rosette-like structures following *Phd2* deletion in the severe *Crb1Crb2*-cKO model. Immunofluorescence images of *Crb1Crb2*-control (A, D, G, J, M, P), *Crb1Crb2*-cKO (B, E, H, K, N, Q), and *Phd2Crb1Crb2*-cKO (C, F, I, L, O, R) retinal sections at P15 (P15; A–C, G–I, M–O) and 1 month of age (1M; D–F, J–L, P–R), stained for rhodopsin (A–F), red/green opsin (G–L), or recoverin (M–R), with DAPI counterstaining (nuclei, gray). In *Crb1Crb2*-control retinas, photoreceptor somata are confined to the ONL and rod and cone outer segment signals are restricted to the apical retina, indicating normal lamination. In *Crb1Crb2*-cKO retinas, rhodopsin-, recoverin-, and red/green opsin-positive cells are dispersed throughout a fused nuclear layer with loss of clear ONL–INL separation. In *Phd2Crb1Crb2*-cKO retinas, these photoreceptor markers are concentrated in rosette-like ONL structures and extend into the rosette lumen. ONL, outer nuclear layer; INL, inner nuclear layer; NL, merged nuclear layer; GCL, ganglion cell layer. Scale bars: 25 µm.

In *Crb1Crb2*-cKO retinas, the ONL and INL were amalgamated into a single, broadened nuclear layer, and rhodopsin-, recoverin-, and red/green opsin-positive cells were diffusely distributed throughout this fused nuclear layer, indicating severe disruption of outer retinal lamination (Fig. 6a B,E,H,K,N,Q). Recoverin-positive photoreceptor somata were also ectopically positioned near or within the ganglion cell layer, demonstrating displacement of photoreceptors into the inner retina (Fig. 6a N,Q). In contrast, *Phd2Crb1Crb2*-cKO retinas displayed rhodopsin-, recoverin-, and red/green opsin-positive photoreceptors arranged in circular profiles corresponding to rosette-like ONL structures at both P15 and 1M (Fig. 6a C,F,I,L,O,R). Nevertheless, some displaced recoverin-positive photoreceptor somata remained detectable near the ganglion cell layer, indicating that *Phd2* deletion did not completely restore retinal lamination (Fig. 6a O,R). In these rosettes, photoreceptor nuclei formed the rosette wall, and rhodopsin and cone opsin signals were present within and oriented toward the rosette lumen, consistent with outer segment–like processes projecting into the lumen (Fig. 6a C,F,I,L). These findings demonstrate that the rosette-like structures in the rescue genotype are composed of rods and cones, that photoreceptor molecular identity is preserved, and that photoreceptor compartments remain spatially organized relative to the rosette lumen in *Phd2Crb1Crb2*-cKO retinas.

### Phd2 deletion promotes local organization of polarity- and synapse-associated proteins within retinal rosettes in the severe Crb1Crb2-cKO-LCA model

Immunohistochemistry for the polarity scaffold protein PALS1, the photoreceptor synaptic scaffold PSD95, the Müller glia/RPE nuclear marker SOX9, and the gliosis marker GFAP was performed on *Crb1Crb2*-control, *Crb1Crb2*-cKO, and *Phd2Crb1Crb2*-cKO rescue retinas at P15 and 1 month (1M) (Fig. 6b). In controls, PALS1 formed a continuous band at the outer limiting membrane (Fig. 6b A,D), PSD95 was confined to a discrete outer plexiform layer (Fig. 6b M,P), SOX9-positive Müller glial cell nuclei formed a regular band in the INL at 1M (Fig. 6b S), and GFAP expression was low and restricted to inner retinal astrocytes and Müller glial cell endfeet within the ganglion cell layer (Fig. 6b G,J), consistent with intact apicobasal polarity, synaptic lamination, and minimal gliosis.

**Figure 6B:**
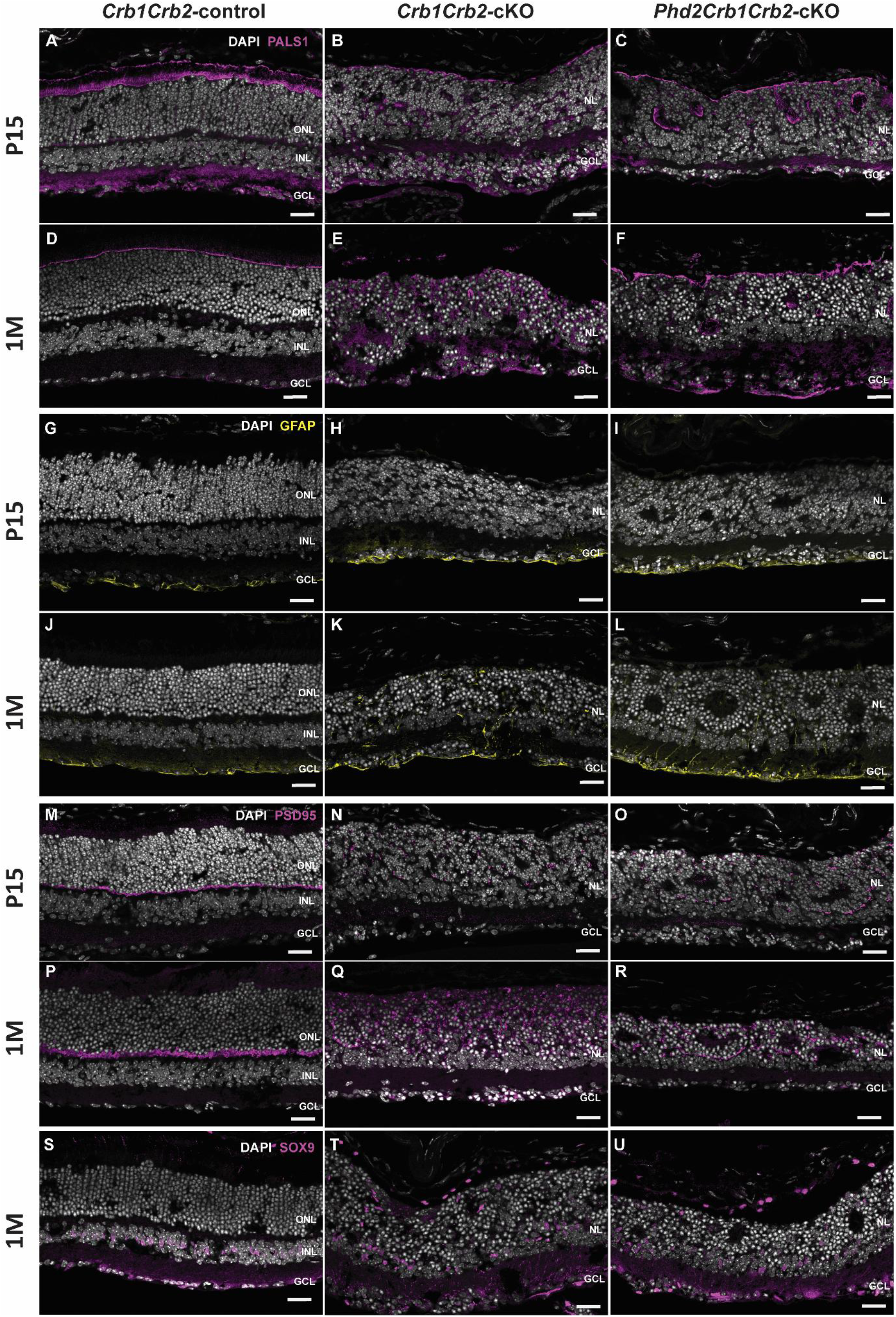
*Phd2* deletion promotes local polarity-associated and synaptic compartmentalization within retinal rosettes in severe *Crb1*/*Crb2* mutants. Representative immunohistochemistry images of *Crb1Crb2-*control (A, D, G, J, M, P, S), *Crb1Crb2-*cKO (B, E, H, K, N, Q, T), and *Phd2Crb1Crb2-*cKO (C, F, I, L, O, R, U) retinas at postnatal day 15 (P15; A–C, G–I, M–O) and 1 month of age (1M; D–F, J– L, P–U). Retinal sections were stained for PALS1 (A–F), GFAP (G–L), PSD95 (M–R), or SOX9 (S–U), with DAPI counterstaining. In controls, PALS1 formed a continuous band at the outer limiting membrane, PSD95 delineated a discrete outer plexiform layer, SOX9-positive Müller glial nuclei were arranged within the INL, and GFAP immunoreactivity was low and restricted mainly to the inner retinal surface. In *Crb1Crb2*-cKO retinas, PALS1 and PSD95 staining was fragmented and dispersed within the merged nuclear layer (NL), while SOX9-positive Müller glial nuclei were displaced into the NL. In *Phd2Crb1Crb2*-cKO retinas, PALS1 outlined rosette-like ONL structures and their lumens, and PSD95 was concentrated along the basal side of the rosettes. Displaced SOX9-positive Müller glial nuclei and GFAP-positive processes remained present. ONL, outer nuclear layer; INL, inner nuclear layer; NL, merged nuclear layer; GCL, ganglion cell layer; OPL, outer plexiform layer; OLM, outer limiting membrane. Scale bars: 25 µm.

In *Crb1Crb2*-cKO retinas, PALS1 and PSD95 labeling became fragmented and dispersed within a merged nuclear layer (NL) (Fig. 6b B,E,N,Q), and SOX9-positive Müller glial cell nuclei were displaced into this nuclear layer (Fig. 6b T), indicating loss of continuous Crumbs polarity, collapse of the synaptic layers, and displacement rather than loss of Müller glial cells. In *Phd2Crb1Crb2*-cKO retinas, PALS1 outlined rosette-like ONL structures and their lumens (Fig. 6b C,F), while PSD95 was concentrated along the basal side of rosettes and in a quasi-continuous band at the mixed nuclear layer level (Fig. 6b O,R), supporting partial restoration of apical junctional polarity and synaptic compartmentalization within the dysplastic outer retina. SOX9-positive Müller glial cell nuclei remained abundant in the nuclear layer at 1M in both mutant genotypes (Fig. 6b T,U). Compared with *Crb1Crb2*-cKO retinas (Fig. 6b H,K), GFAP-positive Müller glial cell processes in *Phd2Crb1Crb2*-cKO retinas appeared slightly more confined to the inner retina and proximal nuclear layer, with no obvious infiltration into rosette cores (Fig. 6b I,L), suggesting that *Phd2* deletion improves outer retinal organization without further exacerbating Müller glial cell gliosis.

To further characterize inflammatory and vascular organization within the dysplastic retina, sections were stained for ionized calcium-binding adaptor molecule 1 (IBA-1) and cluster of differentiation 31 (CD31) (Supplementary Fig. S1). In *Crb1Crb2*-control retinas, IBA-1-positive microglia showed a predominantly laminated inner-retinal distribution at P15 and 1 month. In both *Crb1Crb2*-cKO and *Phd2Crb1Crb2*-cKO retinas, IBA-1-positive profiles were distributed throughout the disorganized retina, including the merged nuclear layer, with no evident qualitative normalization following additional *Phd2* deletion (Supplementary Fig. S1A–F). At 1 month, CD31-positive vascular profiles showed an organized inner-retinal distribution in controls but were irregularly distributed within the disrupted retinal architecture of both mutant genotypes, with more prominent CD31 immunoreactivity observed in *Phd2Crb1Crb2*-cKO retinas (Supplementary Fig. S1G–I). These observations are consistent with the persistent vascular abnormalities detected by fluorescein angiography and indicate that the improved organization of photoreceptor-containing rosettes following *Phd2* deletion was not accompanied by evident normalization of microglial distribution or retinal vascular organization, and prompted subsequent assessment of *Vegfa* transcript expression.

### Phd2 deletion is associated with increased GLUT1 and LDHA protein abundance in the severe Crb1Crb2-cKO retina

To determine whether the structural and functional improvements associated with *Phd2* deletion were accompanied by changes in retinal glucose metabolism, we examined the abundance of proteins involved in glucose uptake and glycolytic metabolism in retinal lysates collected at 1 and 2 months of age. Western blotting was performed for GLUT1, LDHA, and PDK1, and protein abundance was normalized to GAPDH.

At 1 month, GLUT1 abundance did not differ significantly between *Phd2Crb1Crb2*-cKO and *Crb1Crb2*-cKO retinas (Fig. 7A,C). Similarly, LDHA abundance did not differ significantly between the groups at this age (Fig. 7A,D). Thus, the partial structural and functional rescue observed at 1 month was not associated with a statistically significant change in GLUT1 or LDHA protein abundance.

**Figure 7.**
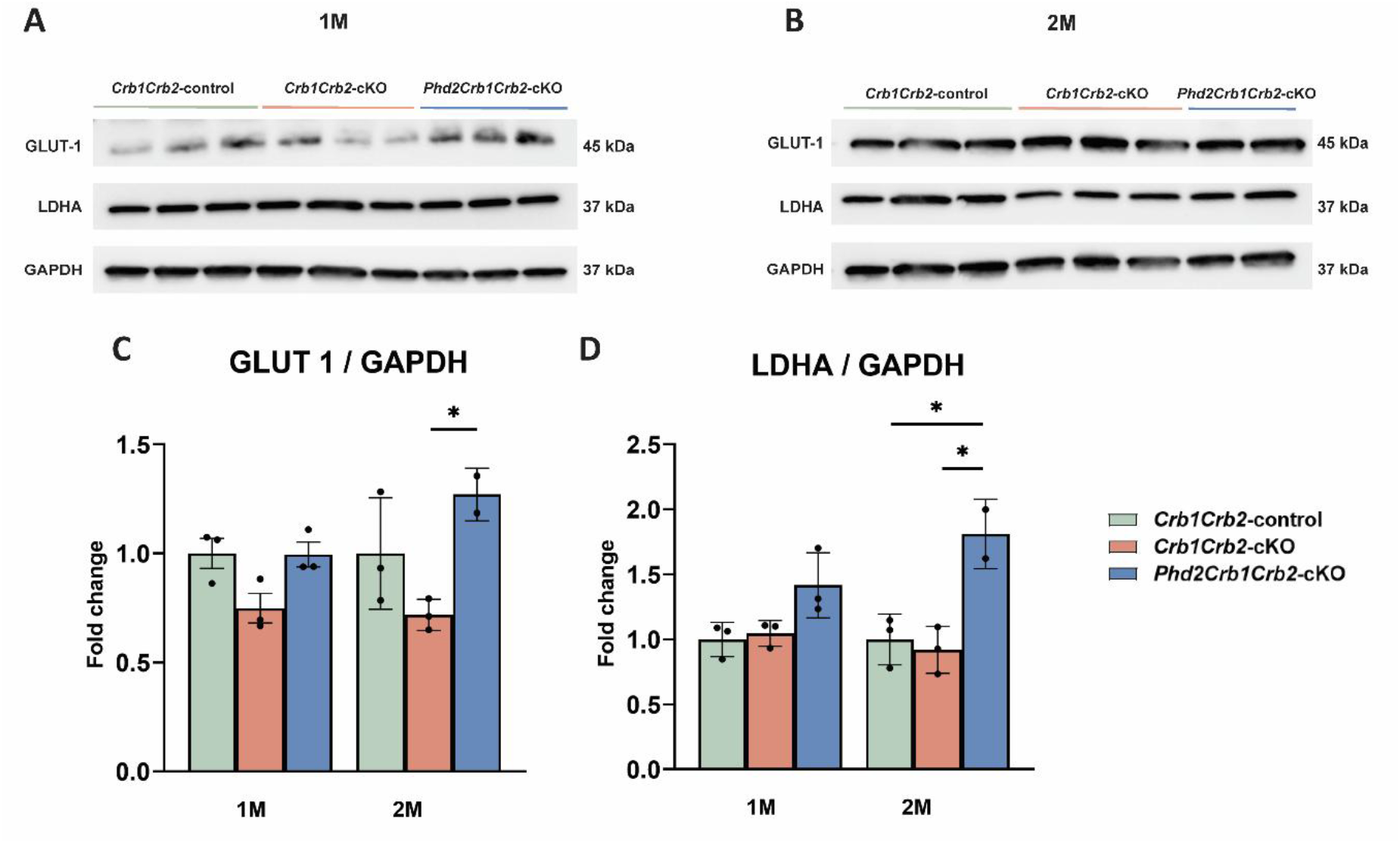
*Phd2* deletion increases retinal GLUT1 and LDHA protein abundance at two months of age. Representative Western blots of retinal lysates from *Crb1Crb2-*control, *Crb1Crb2-*cKO, and *Phd2Crb1Crb2-*cKO mice at (A) 1 month and (B) 2 months of age. Retinal lysates were immunoblotted for GLUT1 (glucose transporter 1), LDHA (lactate dehydrogenase A), and GAPDH (glyceraldehyde-3-phosphate dehydrogenase), with GAPDH serving as the loading control. (C,D) Densitometric quantification of (C) GLUT1 and (D) LDHA protein abundance normalized to GAPDH and expressed as fold change relative to the age-matched control group. At 1 month, GLUT1 and LDHA showed no significant difference in *Phd2Crb1Crb2-*cKO retinas compared with *Crb1Crb2-*cKO retinas. At 2 months, GLUT1 was significantly increased in *Phd2Crb1Crb2-*cKO retinas compared with *Crb1Crb2-*cKO retinas, whereas LDHA was significantly increased compared with both control and *Crb1Crb2-*cKO retinas. Data are presented as mean ± SEM; individual points represent biological replicates. At 1 month, n = 3 mice per genotype; at 2 months, n = 3 for the control and *Crb1Crb2-*cKO groups and n = 2 for the *Phd2Crb1Crb2-*cKO group. Statistical significance was assessed using one-way ANOVA followed by Tukey’s multiple-comparisons test. *p < 0.05.

By 2 months, GLUT1 abundance was significantly increased in *Phd2Crb1Crb2*-cKO retinas compared with *Crb1Crb2*-cKO retinas (Fig. 7B,C). LDHA abundance was also significantly increased in *Phd2Crb1Crb2*-cKO retinas relative to *Crb1Crb2*-cKO retinas and control retinas (Fig. 7B,D). Thus, at 2 months, *Phd2* deletion was associated with significantly higher GLUT1 and LDHA protein abundance in the *Crb1Crb2*-cKO retina. In contrast to GLUT1 and LDHA, PDK1 abundance remained unchanged among control, *Crb1Crb2-*cKO, and *Phd2Crb1Crb2-*cKO retinas at both ages (Supplementary Fig. S2).

### Phd2 deletion induces an early transcriptional response associated with glucose transport and HIF signaling

To determine whether the changes in glycolysis-associated protein abundance following *Phd2* deletion were accompanied by transcriptional regulation, we assessed retinal expression of genes associated with glucose transport, glycolysis, and hypoxia-responsive signaling at 1 and 2 months of age. Relative expression of *Glut1*, *Vegfa*, *Ldha*, *Phd2*, and *Hif1a* was quantified by RT-qPCR in *Crb1Crb2-*cKO phenotypic and *Phd2Crb1Crb2-*cKO rescue retinas (Fig. 8).

**Figure 8.**
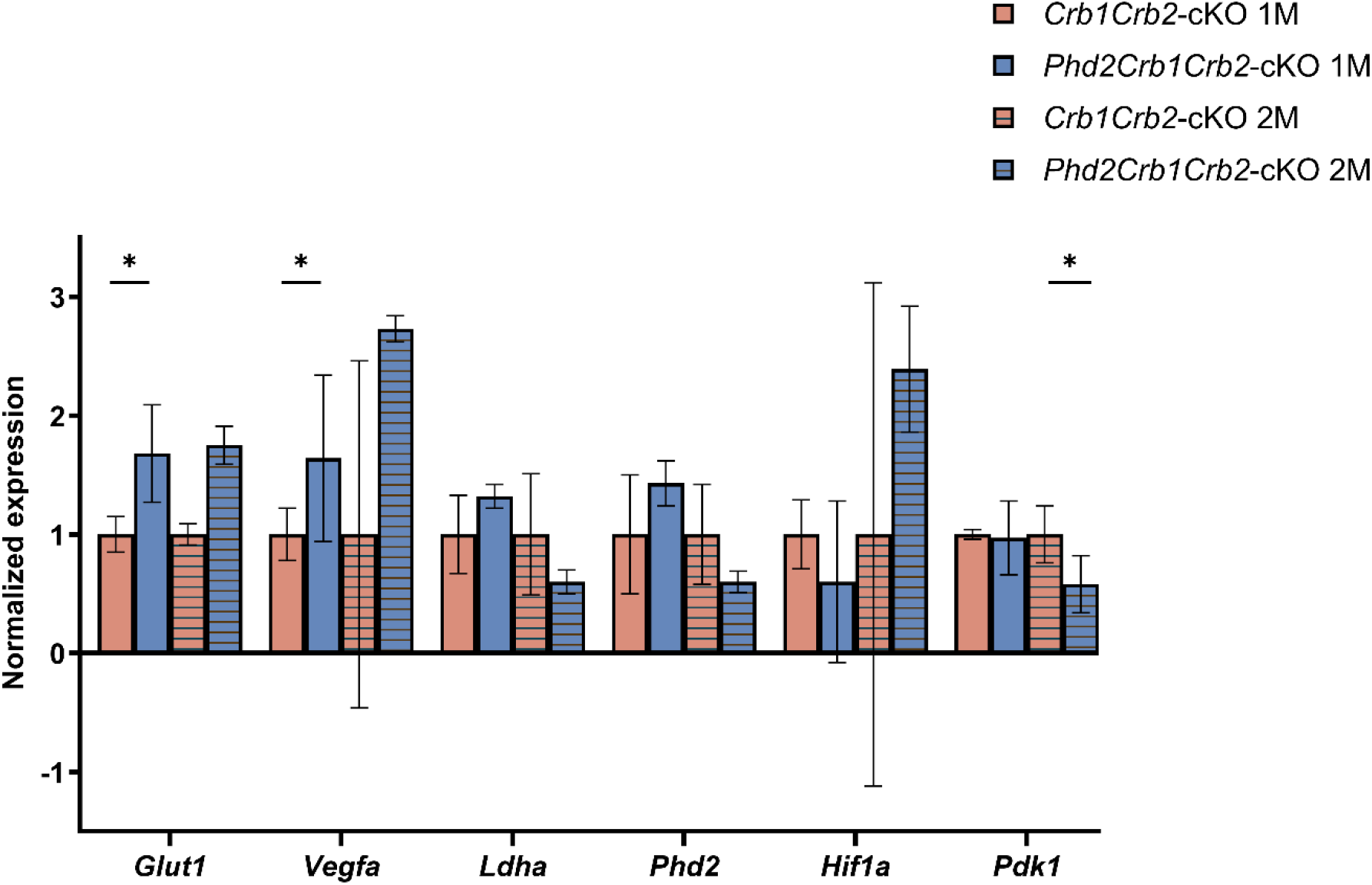
*Phd2* deletion induces early transcriptional activation of glycolytic and hypoxia-associated genes in the severe *Crb1Crb2-*cKO retina. Relative retinal mRNA expression of *Glut1* (glucose transporter 1)*, Vegfa* (vascular endothelial growth factor A)*, Ldha* (lactate dehydrogenase A)*, Phd2 (prolyl hydroxylase domain protein 2), Hif1a (hypoxia-inducible factor 1 alpha), and Pdk1* (pyruvate dehydrogenase kinase 1) in *Crb1Crb2-*cKO and *Phd2Crb1Crb2-*cKO retinas at 1 and 2 months of age. Expression was determined by RT-qPCR, normalized to *Hprt* and *Ef1a*, and expressed as fold change relative to the age-matched *Crb1Crb2*-cKO group. At 1 month, *Glut1* and *Vegfa* expression was significantly increased in *Phd2Crb1Crb2*-cKO retinas compared with *Crb1Crb2*-cKO retinas. At 2 months, *Pdk1* expression was significantly reduced in *Phd2Crb1Crb2*-cKO retinas. No significant differences were detected for *Ldha, Phd2*, or *Hif1a* at either age. Data are mean ± SEM. At 1 month, n= 3 eyes per genotype; at 2 months n=2 eyes per genotype. Statistical comparisons were performed by unpaired two-tailed Welch’s *t*-test. \**p* < 0.05.

At 1 month, *Glut1* and *Vegfa* expression were significantly increased in *Phd2Crb1Crb2-*cKO retinas compared with *Crb1Crb2-*cKO retinas (Fig. 8). At 2 months, *Glut1* and *Vegfa* expression did not differ significantly between *Phd2Crb1Crb2*-cKO and *Crb1Crb2*-cKO retinas. The increase in *Vegfa* transcript abundance was consistent with the more prominent CD31 immunoreactivity observed in *Phd2Crb1Crb2*-cKO retinal sections and coincided with the abnormal retinal vascular phenotype detected by fluorescein angiography (Fig. 3; Supplementary Fig. S1).

*Ldha, Hif1a,* and *Phd2* transcript abundance did not differ significantly between *Phd2Crb1Crb2-cKO* and *Crb1Crb2-*cKO retinas at either age (Fig. 8).

*Pdk1* expression did not differ significantly between groups at 1 month but was significantly lower in *Phd2Crb1Crb2*-cKO retinas at 2 months (Fig. 8). In contrast, PDK1 protein abundance did not differ significantly between groups at either age (Supplementary Fig. S2).

Collectively, these data indicate that *Phd2* deletion is associated with selective alterations in metabolism-associated markers in severe *Crb1Crb2*-cKO retinas. Increased *Glut1* and *Vegfa* transcript levels were observed at 1M, whereas increased GLUT1 and LDHA protein abundance was observed at 2M. In contrast, *Pdk1* transcript levels were significantly reduced at 2 months, while PDK1 protein abundance was unchanged at both time points, indicating selective rather than uniform modulation of glycolysis-associated pathways.

## Discussion

In this study, we show that conditional deletion of *Phd2* in the neural retina partially mitigates the severe retinal phenotype caused by combined Crb1/Crb2 loss. *Phd2* deletion improved rod-driven retinal signaling and visually guided behavior, promoted a more organized rosette-like outer retinal architecture, and elicited temporally dynamic changes in glycolysis- and HIF-associated gene and protein expression. These findings support metabolic reprogramming through modulation of the PHD2/HIF axis as a disease-modifying strategy in a severe developmental model of *CRB1*-associated LCA.

Previous studies have established metabolic reprogramming of photoreceptors through modulation of the PHD2/HIF axis as a potentially mutation-independent strategy for retinal degeneration. Rod-specific disruption of the PHD2/VHL–HIF regulatory pathway enhances aerobic glycolysis, increases expression of glycolytic mediators including GLUT1, LDHA, and PDK1, and preserves photoreceptor structure and function in preclinical models of retinitis pigmentosa (23). Importantly, these studies were performed primarily in models in which retinal lamination is initially established and degeneration is driven by progressive photoreceptor dysfunction or loss. In contrast, the *Crb1Crb2-*cKO retina develops profound abnormalities during retinal development, including disrupted apicobasal polarity, defective lamination, intermingling of nuclear layers, and early functional impairment. The partial structural and functional improvement observed following conditional *Phd2* deletion therefore extends the potential relevance of metabolic reprogramming to a severe developmental model of *CRB1*-associated LCA. However, the incomplete rescue indicates that altered metabolic state may modify the consequences of Crumbs-complex deficiency but cannot fully correct the primary developmental defects resulting from *Crb1* and *Crb2* loss (6, 12).

In the adult mouse retina, CRB1 protein is expressed predominantly in Müller glial cells, whereas CRB2 is expressed in both Müller glial cells and photoreceptors. Thus, combined *Crb1*/*Crb2* deficiency affects multiple retinal cell populations. As Chx10-*Cre* is active broadly in retinal progenitors, conditional *Phd2* deletion may modulate the PHD2/HIF pathway across several neural-retinal cell types (36). However, PHD2 protein abundance and its cell-type-specific distribution were not assessed in the present study, because repeated attempts to detect endogenous PHD2 in retinal lysates by Western blotting yielded no reliable signal, consistent with its reported rapid degradation and consequent difficulty in detection *in vivo* (23). The relative contribution of PHD2/HIF modulation in photoreceptors, Müller glia, and other retinal cell populations to the observed rescue therefore remains unclear. Cell-type-resolved studies will be required to define the cellular basis of this effect.

The functional and structural findings indicate that *Phd2* deletion modifies, but does not normalize, the severe *Crb1Crb2-*cKO phenotype. The increased scotopic a-wave amplitude and reduced b/a-wave ratio suggest improved photoreceptor-driven retinal activity in rescue mice, and the improved optomotor responses indicate that this benefit extends to visually guided behavior. However, scotopic b-wave, photopic b-wave, and flicker responses remained markedly reduced, demonstrating that *Phd2* deletion did not restore normal transmission through the outer retinal circuit or cone-mediated retinal function. Although the outer retina remained disorganized, *Phd2Crb1Crb2-*cKO retinas displayed more clearly delineated rosette-like photoreceptor structures than phenotypic retinas, with rod- and cone-marker-positive cells organized around the rosette lumen. PALS1 and PSD95 labeling further suggested partial re-establishment of local polarity-associated and synaptic compartmentalization within these structures. Because the Crumbs complex is central to photoreceptor–Müller glial cell adhesion, apicobasal polarity, and laminar organization, these findings suggest that metabolic reprogramming may improve the capacity of dysplastic photoreceptors to maintain local tissue organization despite persistent loss of the primary CRB1/CRB2 polarity machinery (37). Consistent with this localized structural effect, IBA-1-positive microglia remained distributed throughout the disorganized retina in both mutant genotypes, suggesting that the improved organization of photoreceptor-containing rosettes was not accompanied by an evident redistribution of retinal microglia (Supplementary Fig. S1).

The molecular findings are consistent with engagement of the metabolic-reprogramming pathway following *Phd2* deletion and provide a potential molecular context for the structural and functional phenotype (38). At 1 month, *Phd2* deletion was associated with significantly increased *Glut1* and *Vegfa* expression, whereas by 2 months GLUT1 and LDHA protein abundance was increased in rescue compared with phenotypic retinas. This temporal pattern is compatible with an early transcriptional response followed by delayed changes at the protein level. GLUT1 facilitates cellular glucose uptake (16), while LDHA supports the conversion of pyruvate to lactate and thereby contributes to the aerobic glycolytic metabolism on which photoreceptors depend for energy production and biosynthetic demands (39). The delayed increase in these proteins is consistent with altered glucose-handling and glycolysis-associated capacity in the rescue retina. However, because these measurements were obtained from whole-retina lysates, the differences may reflect both cell-intrinsic regulation of metabolic proteins and changes in the relative abundance or preservation of the retinal cell populations expressing them. Moreover, because glucose uptake, lactate production, and metabolic flux were not measured directly, these findings provide evidence of altered glycolysis-associated gene and protein expression rather than direct proof of increased glycolytic flux. The absence of a corresponding change in PDK1 indicates that the metabolic response to *Phd2* deletion did not uniformly involve all glycolysis-associated proteins examined. Furthermore, the absence of a detectable reduction in *Phd2* transcript abundance may reflect mosaic *Chx10*-Cre-mediated recombination within the neural retina, together with detection of transcripts from unrecombined cells by primers that do not specifically distinguish the recombined *Phd2* allele.

The early induction of *Glut1* and *Vegfa* is consistent with engagement of hypoxia-responsive signaling following conditional *Phd2* deletion. In particular, increased *Vegfa* expression raises an important consideration for PHD2-directed therapeutic strategies. VEGFA is a canonical HIF-responsive factor with essential functions in retinal vascular development, tissue perfusion, and neuronal homeostasis (40). However, excess VEGFA signaling can disrupt blood–retinal barrier integrity and promote vascular permeability (41). In the present study, elevated *Vegfa* expression at 1 month coincided with persistent vascular abnormalities and apparent fluorescein leakage observed by fluorescein angiography (Fig. 3). Histologically, CD31-positive vascular profiles were irregularly distributed in both mutant genotypes, with more prominent CD31 immunoreactivity observed in *Phd2Crb1Crb2*-cKO retinas (Supplementary Fig. S1). Together, these findings are consistent with a vascular response accompanying the early increase in *Vegfa* expression following Phd2 deletion (Fig. 3). Although these findings do not establish that VEGFA induction directly causes the angiographic phenotype, they suggest that PHD2/HIF modulation may have both potentially adaptive metabolic effects and vascular consequences. Further analysis of *Phd2* ablation in another Crumbs-deficient background, such as *Phd2Crb1^(+/−)^Crb2*-cKO mice, may help distinguish vascular alterations associated with PHD2/HIF pathway modulation from those secondary to the profound retinal dysplasia in the severe *Crb1Crb2*-cKO model. Future studies should identify the retinal cell type(s) responsible for VEGFA expression and directly quantify vascular permeability, retinal blood flow, and blood–retinal barrier integrity following *Phd2* deletion.

Several limitations should be considered when interpreting the present findings. Molecular analyses were performed using whole-retina lysates and therefore cannot assign changes in glycolysis- and hypoxia-associated gene and protein expression to specific retinal cell populations. Importantly, differences in bulk protein abundance may also partly reflect genotype- and age-dependent differences in retinal cellular composition rather than changes in protein expression within individual cells. This consideration is particularly relevant in the severely dysplastic *Crb1Crb2-*cKO retina, where progressive alterations in retinal architecture and the relative abundance or preservation of different retinal cell populations may substantially influence the composition of whole-retina lysates. Thus, the higher GLUT1 and LDHA abundance detected in *Phd2*-deleted retinas at two months could reflect a combination of cell-intrinsic metabolic regulation and differences in the cellular composition or preservation of the retina. Cell-type-resolved approaches and direct metabolic assays will be required to define how *Phd2* deletion alters retinal metabolism.

From a translational perspective, metabolic reprogramming may offer an advantage over mutation-specific interventions by targeting shared cellular stress pathways across genetically diverse retinal dystrophies. In the present developmental *CRB1*-LCA model, *Phd2* deletion did not correct the underlying Crumbs-complex polarity defect but partially improved retinal structure and function, suggesting that modulation of the PHD2/HIF axis may enhance the metabolic resilience of cells exposed to chronic architectural and metabolic stress. PHD2/HIF modulation may therefore be most appropriately considered as a disease-modifying strategy or as a potential complement to gene-specific approaches, rather than as a stand-alone correction for severe *CRB1*-associated disease. Future work should define the optimal therapeutic window, safety, target cell population, and degree of PHD2/HIF modulation required to maximize photoreceptor benefit while limiting adverse vascular effects.

## Conclusion

Conditional retinal *Phd2* deletion partially ameliorated visual dysfunction and structural disorganization in the severe *Crb1Crb2-*cKO model of *CRB1*-associated LCA. The improvement in rod-driven function, visually guided behavior, and photoreceptor organization was accompanied by temporally dynamic changes in glycolysis- and HIF-associated gene and protein expression, including increased *Glut1* and *Vegfa* expression at 1 month and elevated GLUT1 and LDHA protein abundance at 2 months. Although *Phd2* deletion did not restore normal retinal lamination or fully recover retinal function, these findings support PHD2/HIF-mediated metabolic reprogramming as a potential disease-modifying strategy for severe *CRB1*-associated retinal degeneration.

## Materials and methods

### Animals

All procedures concerning animals were carried out in accordance with the E.U. Directive 2010/63/EU for animal experiments and with permission from the Dutch Central Authority for Scientific Procedures on Animals (CCD), permit number AVD11600202317512. All mice used were maintained with a 99.9% C57BL/6J genetic background. Animals were kept in a 12 hours day/night cycle and supplied with food and water ad libitum. Mice did not have *rd8* or *pde6b* mutations. For electroretinography (ERG), micedark-adapted overnight before recording.

To generate the three experimental genotypes as littermates, we crossed *Phd2*^F/+^*Crb1*^−/−^*Crb2*^F/F^Chx*10Cre*^Tg/+^ mice with *Phd2*^F/+^*Crb1*^−/−^*Crb2*^F/F^ mice. The Chx10-*Cre* transgene drives Cre recombinase expression throughout the developing neural retina from embryonic day (E)11.5, in all retinal cell types except the retinal pigment epithelium (27) as previously described (28). This cross yielded, within the same litters: (i) phenotypic controls (*Crb1*^−/−^*Crb2*^F/F^, lacking Chx*10*-*Cre*; here abbreviated as *Crb1Crb2-*control), in which *Crb2* remains intact and only the mild *Crb1*-null phenotype is manifest; (ii) phenotypic (severe LCA) mice (*Crb1*^−/−^*Crb2*^F/F^Chx10*Cre*^Tg/+^, *Phd2*^+/+^; here abbreviated as *Crb1Crb2-*cKO), in which conditional deletion of *Crb2* on the *Crb1*-null background abolishes *Crb1*/*Crb2* redundancy and reproduces the severe early-onset LCA phenotype previously characterized (11); and (iii) *Phd2*-rescue mice (*Phd2*^F/F^*Crb1*^−/−^*Crb2*^F/F^Chx10*Cre*^Tg/+^; here abbreviated as *Phd2Crb1Crb2-*cKO), in which both *Crb2* and *Phd2* are concurrently deleted throughout the neural retina.

Ear biopsies were genotyped commercially by Transnetyx (Cordova, TN, USA) using real-time PCR with strain-specific probes. Mice were genotyped for *Crb1*, *Crb2*, *Phd2* and *Cre* alleles, and only animals confirmed to carry the desired genotypes were used for subsequent experiments. Male and female mice were used in all groups.

### Electroretinography (ERG)

Dark- and light-adapted ERGs were recorded under dim red light using an Espion E2 system (Diagnosys, LLC, Lowell, MA, USA). Recordings were obtained from 1-month-old (1M) *Crb1Crb2-*control*, Crb1Crb2-*cKO and *Phd2Crb1Crb2-*cKO mice. Mice were anesthetized by intraperitoneal injection of ketamine (100 mg kg⁻¹) and xylazine (10 mg kg⁻¹), and pupils were dilated with tropicamide (5 mg ml⁻¹). Body temperature was maintained using a temperature-controlled heating pad. Reference and ground platinum electrodes were placed subcutaneously in the scalp and at the base of the tail, respectively, and ERGs were recorded from both eyes using gold wire electrodes. Hypromellose eye drops (3 mg ml⁻¹; Teva, The Netherlands) were applied between recordings to prevent corneal drying. Single-flash scotopic and photopic ERGs were elicited using white light (6500 K). Signals were band-pass filtered at 0.3– 300 Hz. Scotopic responses were recorded following dark adaptation at flash intensities of −4, −3, −2, −1, 0, 1, 1.5, and 1.9 log cd·s m⁻². Photopic responses were recorded after 10 min of light adaptation to a background of 30 cd m⁻², using flash intensities of −2, −1, 0, 1, 1.5, and 1.9 log cd·s m⁻². The following numbers of mice were used per time point: 1-month-old: *Crb1Crb2-*control *(n=*4)*, Crb1Crb2-*cKO (n=6) and *Phd2Crb1Crb2-*cKO (n=5). Statistical analysis of ERG amplitudes was performed in GraphPad Prism (version 10.6.1; GraphPad Software, San Diego, CA) using one-way ANOVA at each time point to compare responses between experimental groups. When the ANOVA indicated a significant effect, pairwise differences between all groups were assessed using Tukey’s multiple comparisons post hoc test, with a family-wise significance level of α = 0.05. Data are presented as mean ± SEM, and multiplicity-adjusted *P* values from the post hoc tests are reported in the figure legends.

### Optokinetic Head Tracking Reflex (OKT)

Spatial frequency and contrast sensitivity were assessed using an OptoMotry system (Cerebral Mechanics, Lethbridge, AB, Canada). One and two month old *Crb1Crb2-*control*, Crb1Crb2-*cKO and *Phd2Crb1Crb2-*cKO mice were placed on a central platform surrounded by four computer monitors forming a virtual cylinder displaying a rotating vertical sine wave grating (12° s⁻¹), as previously described. Tracking of the stimulus by reflexive head movements was scored as a positive response.

Spatial frequency thresholds were determined using a staircase paradigm beginning at 0.042 cycles deg⁻¹ with 100% contrast. Contrast sensitivity thresholds were assessed at three spatial frequencies (0.031, 0.064 and 0.092cycles deg⁻¹), and contrast sensitivity was calculated as the inverse of the contrast threshold for each frequency.

The numbers of mice used for the optokinetic head tracking reflex measurements per time point were as follows: 1-month-old: *Crb1Crb2-*control *(n=*13)*, Crb1Crb2-*cKO (n=14) and *Phd2Crb1Crb2-*cKO (n=14). Statistical significance was calculated by using one-way ANOVA followed by Tukey’s multiple-comparisons test.

### Confocal scanning laser ophthalmoscopy (cSLO), fluorescein angiography (FA) and spectral-domain optical coherence tomography (SD-OCT)

Retinal morphology of anesthetized animals was visualized using confocal scanning laser ophthalmoscopy (cSLO) with Heidelberg Retina Angiograph 2 system (HRA2; Heidelberg Engineering, Heidelberg, Germany), according to previously described procedures (42). Fluorescein angiography

(FA) was carried out following injection of 200mg/kg fluorescein sodium to visualize retinal vasculature and assess blood–retinal barrier integrity (34). Spectral-domain optical coherence tomography (SD-OCT) imaging was performed with a EnvisuTM R2210 VHR (Leica, USA) to obtain cross-sectional retinal scans using a broadband superluminescent diode at λ = 870 nm as low-coherence light source.

### Morphological and Immunohistochemical Analysis

Eyes from P15 and 1-month-old (1M) old *Crb1Crb2-*control*, Crb1Crb2-*cKO and *Phd2Crb1Crb2-*cKO mice were collected (*n* = 3-5 eyes/age/group). Enucleated eyes were fixed in 4% paraformaldehyde (PFA) in phosphate-buffered saline (PBS) for 20 min at room temperature and rinsed in PBS. Samples were dehydrated through a graded ethanol series (30%, 50%, and 70% for 30 min each; 90% and 100% ethanol twice for 30 min each), followed by incubation in a 1:1 mixture of 100% ethanol and Technovit 7100 (Kulzer) for 30 min. Tissues were then infiltrated overnight at 4 °C in Technovit 7100 supplemented with Hardener I (1 g per 100 ml; Technovit 1). Samples were embedded the following day in Technovit 1 containing Hardener II (15:1; Technovit 2), oriented, and polymerized overnight. Polymerized blocks were reinforced with Technovit 3040 (Kulzer) mixed with universal hardener (5 scoops per 10 ml) and allowed to harden for several hours. Blocks were sectioned at 3 µm using a microtome. Sections were air-dried, counterstained with 0.1% toluidine blue, and mounted with Entellan (Sigma-Aldrich) under glass coverslips. Whole-slide images were acquired using a Pannoramic 250 scanner (3DHISTECH), and images were visualized and processed using SlideViewer (v2.9).

For immunohistochemical analysis, eyes were enucleated and fixed for 20 min in 4% paraformaldehyde in PBS. Subsequently, the tissues were cryo-protected with 15% and 30% sucrose in PBS, embedded in Tissue-Tek O.C.T. Compound (Sakura, Finetek, Alphen aan den Rijn, The Netherlands), and used for cryosectioning. Cryosections (8 µm) were rehydrated in PBS and blocked for 1 h at room temperature in pre-incubation buffer consisting of 10% normal goat serum (Thermo Fisher Scientific, Waltham, MA, USA), 0.4% Triton X-100 and 1% bovine serum albumin in PBS. Sections were then incubated overnight at 4 °C with the primary antibodies in PBS containing 0.3% normal goat serum, 0.4% Triton X-100 and 1% bovine serum albumin. After two 15-min washes in PBS, sections were incubated for 1 h at room temperature with fluorescently labeled secondary antibody [goat anti-rabbit IgGs conjugated to Alexa 555 (1:1000; Abcam, Cambridge, UK)] diluted in PBS containing 1% BSA. Details of all primary antibodies used are provided in Supplementary Table S1. Following two additional 15-min PBS washes, slides were mounted with Vectashield Antifade Mounting Medium with DAPI (Vector Laboratories, Newark, CA, USA) for nuclear counterstaining. Images were acquired using a Leica TCS SP8 confocal microscope (Leica Microsystems, Wetzlar, Germany) with Leica Application Suite X v3.7.0.20979.

### Transmission Electron Microscopy

Eyes from 1-month-old *Crb1Crb2*-control and *Crb1Crb2*-cKO mice were collected and fixed in 2% paraformaldehyde and 1.5% glutaraldehyde in 0.1 M cacodylate buffer at room temperature for 1 h (n = 1 per group). The eyes were cut into two halves, with the lens located on one side and the optic nerve head (blind spot) on the other, and the vitreous body was removed. Both halves were returned to the fixative for an additional 1 h. After rinsing, the samples were post-fixed in 1% OsO₄/1,5% potassiumferricyanide in 0.1 M cacodylate buffer and rinsed again. Following fixation and post-fixation, both halves were further trimmed so that the optic nerve head or lens/iris was positioned centrally within the respective tissue block.

The samples were dehydrated through an ascending ethanol series, followed by infiltration with mixtures of acetone and EPON (LX112) and subsequently with pure EPON. The pieces of retina ware positioned in molds that way to obtain the most optimal cross sections of the retina. After polymerization of the EPON ultrathin sections (80 nm) were prepared, stained with uranyl acetate and lead citrate. Images were acquired on a digital camera (One View, Gatan Inc., Pleasanton, USA) mounted on a 120 kV transmission electron microscope (Tecnai T12 with a twin objective lens, Thermofisher, Eindhoven, The Netherlands). Overlapping images were collected and stitched together into one image, as previously described (43).

### Quantification of the number of rosettes and GCL thickness

All retinal morphometric analyses were performed on digital whole-slide images using SlideViewer software (version 2.9; 3DHISTECH). For each animal, three representative retinal sections were selected, and in each section the neurosensory retina was manually delineated to define the region of interest. Ganglion cell layer (GCL) thickness was quantified by acquiring five cross-sectional measurements per section at predefined positions across the retina; these values were averaged to obtain a mean thickness per section, which was then averaged across the three sections to yield a mean GCL thickness per animal, and this procedure was applied identically to all control, phenotypic, and rescue groups. Rosette pathology was quantified by identifying structures that met strict morphological criteria, namely a circular or ring-like arrangement of darkly stained nuclei forming a fully closed perimeter, and the total number of such rosettes per section was normalized to the analyzed retinal area and expressed as rosettes per mm² to account for variability in tissue size.

### RNA isolation, cDNA synthesis, and RT-qPCR

Retinas from 1- and 2-month-old *Crb1Crb-*control, *Crb1Crb2-*cKO, and *Phd2Crb1Crb2-*cKO mice were dissected, snap-frozen, and stored at −80 °C until total RNA was isolated using the RNeasy Mini Kit (Qiagen, Hilden, Germany; cat. no. 74104). Retinas were manually homogenized in lysis buffer by repeated pipetting, and RNA concentration and purity were determined using a NanoDrop spectrophotometer (Thermo Fisher Scientific). Samples were stored at −80 °C until cDNA synthesis. cDNA was synthesized from 500 ng total RNA using the TaqMan™ Reverse Transcription Kit (Applied Biosystems; cat. no. N8080234) on a C1000 Touch™ Thermal Cycler (Bio-Rad). cDNA was diluted 1:20 before qPCR, which was performed using iQ™ SYBR® Green Supermix (Bio-Rad; cat. no. 1708880) on a CFX Opus 384 Real-Time PCR System (Bio-Rad). Gene expression was normalized to *Hprt* and *Ef1a1*, and normalized expression was calculated relative to age-matched *Crb1Crb2*-cKO retinas using the 2^−ΔΔ*C*^*^t^* method, with the mean expression of the *Crb1Crb2*-cKO group set to 1. All primer sequences used for qPCR are provided in Supplementary Table S2.

### Western blotting

Retinas from 1- and 2-month-old *Crb1Crb-*control, *Crb1Crb2-*cKO, and *Phd2Crb1Crb2-*cKO mice were dissected, snap-frozen, and stored at −80 °C until protein extraction. Each retina was homogenized in 100 µL ice-cold RIPA buffer supplemented with cOmplete™ protease inhibitor (Roche), incubated on ice for 30 min, and protein concentration was determined using the Pierce™ BCA Protein Assay Kit (Thermo Fisher Scientific; cat. no. 23225). Equal protein amounts (20 µg/lane) were separated on 4–20% Mini-PROTEAN® TGX™ precast gels (Bio-Rad), transferred to PVDF membranes using the Trans-Blot® Turbo™ system (Bio-Rad), and blocked in 5% milk in PBS-T. Membranes were incubated overnight at 4 °C with primary antibodies, followed by HRP-conjugated secondary antibodies for 1 h at room temperature. Details of all primary antibodies used are provided in Supplementary Table S1. Signals were detected using Clarity™ Western ECL Substrate (Bio-Rad) on a ChemiDoc™ Imaging System (Bio-Rad). Band intensities were quantified in Fiji and normalized to GAPDH. For both qPCR and Western blot analyses, *n* = 3 mice per genotype at 1 month and for the control and phenotypic groups at 2 months; the 2-month rescue group comprised *n* = 2 mice.

### Statistical Analysis

Statistical analyses were performed using GraphPad Prism version 10.6.1 (GraphPad Prism, RRID: SCR_002798; GraphPad Software, San Diego, CA). Unless otherwise specified in the figure legends, data are presented as mean ± SEM and comparisons among experimental groups were performed using one-way ANOVA followed by Tukey’s multiple-comparisons test. Alternative statistical tests and methods of data presentation are specified in the corresponding figure legends. Statistically significant values: * *p* < 0.05, ** *p* < 0.01, *** *p* < 0.001, **** *p* < 0.0001 or *ns=* not significant.

## Supporting information

Phd2 loss rescues severe LCA_Supplementary

## Data availability

The supporting data of this study can be obtained from the corresponding author upon reasonable request.

## Author Contributions

Conceptualization, V.K. and J.W.; Methodology, V.K. and J.W.; Formal analysis, V.K. and J.W.; Investigation, V.K., C.A.A. and X.L.; Data curation, V.K.; Writing—original draft preparation, V.K.; Writing—review and editing, V.K. and J.W.; Visualization, V.K.; Supervision, J.W.; Project administration, J.W.; Funding acquisition, J.W.

## Acknowledgments

The authors all team members for valuable discussions and advice, and the students from the RetoRNA doctoral network for critical assessment and exchange of ideas.

## References

1. Huang CH, Yang CM, Yang CH, Hou YC, Chen TC. Leber’s Congenital Amaurosis: Current Concepts of Genotype-Phenotype Correlations. Genes (Basel). 2021;12(8).

2. Daich Varela M, Cabral de Guimaraes TA, Georgiou M, Michaelides M. Leber congenital amaurosis/early-onset severe retinal dystrophy: current management and clinical trials. Br J Ophthalmol. 2022;106(4):445–51.

3. Coussa RG, Lopez Solache I, Koenekoop RK. Leber congenital amaurosis, from darkness to light: An ode to Irene Maumenee. Ophthalmic Genet. 2017;38(1):7–15.

4. Lotery AJ, Jacobson SG, Fishman GA, Weleber RG, Fulton AB, Namperumalsamy P, et al. Mutations in the CRB1 gene cause Leber congenital amaurosis. Arch Ophthalmol. 2001;119(3):415–20.

5. van de Pavert SA, Sanz AS, Aartsen WM, Vos RM, Versteeg I, Beck SC, et al. Crb1 is a determinant of retinal apical Muller glia cell features. Glia. 2007;55(14):1486–97.

6. Alves CH, Pellissier LP, Wijnholds J. The CRB1 and adherens junction complex proteins in retinal development and maintenance. Prog Retin Eye Res. 2014;40:35–52.

7. van Rossum AG, Aartsen WM, Meuleman J, Klooster J, Malysheva A, Versteeg I, et al. Pals1/Mpp5 is required for correct localization of Crb1 at the subapical region in polarized Muller glia cells. Hum Mol Genet. 2006;15(18):2659–72.

8. Mehalow AK, Kameya S, Smith RS, Hawes NL, Denegre JM, Young JA, et al. CRB1 is essential for external limiting membrane integrity and photoreceptor morphogenesis in the mammalian retina. Hum Mol Genet. 2003;12(17):2179–89.

9. Aleman TS, Cideciyan AV, Aguirre GK, Huang WC, Mullins CL, Roman AJ, et al. Human CRB1-associated retinal degeneration: comparison with the rd8 Crb1-mutant mouse model. Invest Ophthalmol Vis Sci. 2011;52(9):6898–910.

10. van de Pavert SA, Kantardzhieva A, Malysheva A, Meuleman J, Versteeg I, Levelt C, et al. Crumbs homologue 1 is required for maintenance of photoreceptor cell polarization and adhesion during light exposure. J Cell Sci. 2004;117(Pt 18):4169–77.

11. Pellissier LP, Lundvig DM, Tanimoto N, Klooster J, Vos RM, Richard F, et al. CRB2 acts as a modifying factor of CRB1-related retinal dystrophies in mice. Hum Mol Genet. 2014;23(14):3759–71.

12. Pellissier LP, Alves CH, Quinn PM, Vos RM, Tanimoto N, Lundvig DM, et al. Targeted ablation of CRB1 and CRB2 in retinal progenitor cells mimics Leber congenital amaurosis. PLoS Genet. 2013;9(12):e1003976.

13. Hanna J, David LA, Touahri Y, Fleming T, Screaton RA, Schuurmans C. Beyond Genetics: The Role of Metabolism in Photoreceptor Survival, Development and Repair. Front Cell Dev Biol. 2022;10:887764.

14. Narayan DS, Chidlow G, Wood JP, Casson RJ. Glucose metabolism in mammalian photoreceptor inner and outer segments. Clin Exp Ophthalmol. 2017;45(7):730–41.

15. Peachey NS, Yu M, Han JYS, Lengacher S, Magistretti PJ, Pellerin L, et al. Impact of MCT1 Haploinsufficiency on the Mouse Retina. Adv Exp Med Biol. 2018;1074:375–80.

16. Daniele LL, Han JYS, Samuels IS, Komirisetty R, Mehta N, McCord JL, et al. Glucose uptake by GLUT1 in photoreceptors is essential for outer segment renewal and rod photoreceptor survival. FASEB J. 2022;36(8):e22428.

17. Hass DT, Giering E, Han JYS, Bisbach CM, Pandey K, Robbings BM, et al. In vivo exchange of glucose and lactate between photoreceptors and the retinal pigment epithelium. eLife. 2024;eLife 105738.

18. Du J, Yanagida A, Knight K, Engel AL, Vo AH, Jankowski C, et al. Reductive carboxylation is a major metabolic pathway in the retinal pigment epithelium. Proc Natl Acad Sci U S A. 2016;113(51):14710–5.

19. Kanow MA, Giarmarco MM, Jankowski CS, Tsantilas K, Engel AL, Du J, et al. Biochemical adaptations of the retina and retinal pigment epithelium support a metabolic ecosystem in the vertebrate eye. Elife. 2017;6.

20. Owen N, Toms M, Tian Y, Toualbi L, Richardson R, Young R, et al. Loss of the crumbs cell polarity complex disrupts epigenetic transcriptional control and cell cycle progression in the developing retina. J Pathol. 2023;259(4):441–54.

21. Adani E, Villa Vasquez SS, Lovino M, Bighinati A, Cappellino L, D’Alessandro S, et al. Modulation of Aerobic Glycolysis Genes During the Progression of Retinitis Pigmentosa. Invest Ophthalmol Vis Sci. 2026;67(1):49.

22. Pan WW, Wubben TJ, Besirli CG. Photoreceptor metabolic reprogramming: current understanding and therapeutic implications. Communications Biology. 2021;4(1).

23. Nolan ND, Cui X, Robbings BM, Demirkol A, Pandey K, Wu WH, et al. CRISPR editing of anti-anemia drug target rescues independent preclinical models of retinitis pigmentosa. Cell Rep Med. 2024;5(4):101459.

24. Zhang L, Du J, Justus S, Hsu CW, Bonet-Ponce L, Wu WH, et al. Reprogramming metabolism by targeting sirtuin 6 attenuates retinal degeneration. J Clin Invest. 2016;126(12):4659–73.

25. Alves CH, Sanz AS, Park B, Pellissier LP, Tanimoto N, Beck SC, et al. Loss of CRB2 in the mouse retina mimics human retinitis pigmentosa due to mutations in the CRB1 gene. Hum Mol Genet. 2013;22(1):35–50.

26. Takeda K, Ho VC, Takeda H, Duan LJ, Nagy A, Fong GH. Placental but not heart defects are associated with elevated hypoxia-inducible factor alpha levels in mice lacking prolyl hydroxylase domain protein 2. Mol Cell Biol. 2006;26(22):8336–46.

27. Nishida A, Furukawa A, Koike C, Tano Y, Aizawa S, Matsuo I, et al. Otx2 homeobox gene controls retinal photoreceptor cell fate and pineal gland development. Nature Neuroscience. 2003;6(12):1255–63.

28. Ivanova E, Hwang GS, Pan ZH. Characterization of transgenic mouse lines expressing Cre recombinase in the retina. Neuroscience. 2010;165(1):233–43.

29. Wang YV, Weick M, Demb JB. Spectral and temporal sensitivity of cone-mediated responses in mouse retinal ganglion cells. J Neurosci. 2011;31(21):7670–81.

30. Buck TM, Vos RM, Alves CH, Wijnholds J. AAV-CRB2 protects against vision loss in an inducible CRB1 retinitis pigmentosa mouse model. Mol Ther Methods Clin Dev. 2021;20:423–41.

31. Pellissier LP, Quinn PM, Alves CH, Vos RM, Klooster J, Flannery JG, et al. Gene therapy into photoreceptors and Muller glial cells restores retinal structure and function in CRB1 retinitis pigmentosa mouse models. Hum Mol Genet. 2015;24(11):3104–18.

32. Lin V, Lee W, Kang EY, Liu PK, Wang NK. Outer retinal tubulation associated with photoreceptor degeneration. Prog Retin Eye Res. 2026;111:101435.

33. Goldberg NR, Greenberg JP, Laud K, Tsang S, Freund KB. Outer retinal tubulation in degenerative retinal disorders. Retina. 2013;33(9):1871–6.

34. Fischer MD, Huber G, Beck SC, Tanimoto N, Muehlfriedel R, Fahl E, et al. Noninvasive, in vivo assessment of mouse retinal structure using optical coherence tomography. PLoS One. 2009;4(10):e7507.

35. Stuck MW, Conley SM, Naash MI. Defects in the outer limiting membrane are associated with rosette development in the Nrl-/-retina. PLoS One. 2012;7(3):e32484.

36. Rowan S, Cepko CL. Genetic analysis of the homeodomain transcription factor Chx10 in the retina using a novel multifunctional BAC transgenic mouse reporter. Dev Biol. 2004;271(2):388–402.

37. Park B, Alves CH, Lundvig DM, Tanimoto N, Beck SC, Huber G, et al. PALS1 is essential for retinal pigment epithelium structure and neural retina stratification. J Neurosci. 2011;31(47):17230–41.

38. Olivares-Gonzalez L, Martinez-Fernandez de la Camara C, Hervas D, Millan JM, Rodrigo R. HIF-1alpha stabilization reduces retinal degeneration in a mouse model of retinitis pigmentosa. FASEB J. 2018;32(5):2438–51.

39. Rajala A, Bhat MA, Teel K, Gopinadhan Nair GK, Purcell L, Rajala RVS. The function of lactate dehydrogenase A in retinal neurons: implications to retinal degenerative diseases. PNAS Nexus. 2023;2(3):pgad038.

40. Vinores SA, Xiao WH, Aslam S, Shen J, Oshima Y, Nambu H, et al. Implication of the hypoxia response element of the Vegf promoter in mouse models of retinal and choroidal neovascularization, but not retinal vascular development. J Cell Physiol. 2006;206(3):749–58.

41. Roberts WG, Palade GE. Increased microvascular permeability and endothelial fenestration induced by vascular endothelial growth factor. J Cell Sci. 1995;108 ( Pt 6):2369–79.

42. Seeliger MW, Beck SC, Pereyra-Munoz N, Dangel S, Tsai JY, Luhmann UF, et al. In vivo confocal imaging of the retina in animal models using scanning laser ophthalmoscopy. Vision Res. 2005;45(28):3512–9.

43. Faas FG, Avramut MC, van den Berg BM, Mommaas AM, Koster AJ, Ravelli RB. Virtual nanoscopy: generation of ultra-large high resolution electron microscopy maps. J Cell Biol. 2012;198(3):457–69.

