## Supplementary material for "Retinal *Phd2* ablation partially rescues *Crb1Crb2*-associated Leber congenital amaurosis and alters metabolic signaling": Phd2 loss rescues severe LCA_Supplementary

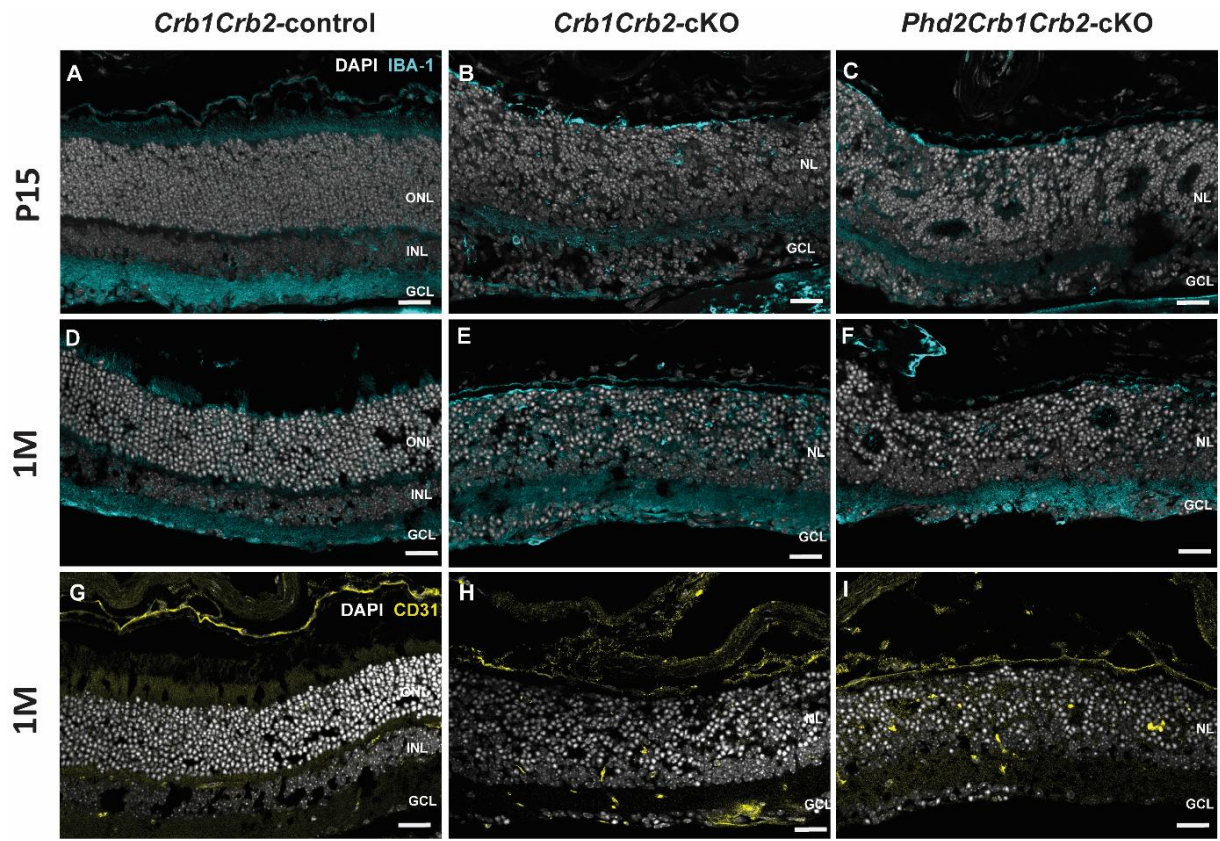

**Supplementary Figure S1. Microglial and vascular organization in *Crb1Crb2*-cKO retinas following *Phd2* deletion.** Representative immunofluorescence images of retinal sections from *Crb1Crb2*-control (A, D, G), *Crb1Crb2*-cKO (B, E, H), and *Phd2Crb1Crb2*-cKO (C, F, I) mice. Sections were stained for IBA1 to visualize microglia at P15 (A–C) and 1M (1M; D–F), or for CD31 to visualize retinal endothelial cells at 1 month (G–I), with DAPI counterstaining of nuclei (gray). In *Crb1Crb2*-control retinas, IBA1-positive profiles showed a laminated inner-retinal distribution, whereas in *Crb1Crb2*-cKO and *Phd2Crb1Crb2*-cKO retinas, IBA1-positive profiles were distributed throughout the disorganized retinal tissue, including the merged nuclear layer. CD31-positive vascular profiles were organized within the inner retina in controls but appeared irregularly distributed within the disrupted retinal architecture of both mutant genotypes. No evident qualitative normalization of microglial distribution or vascular organization was observed following additional *Phd2* deletion. ONL, outer nuclear layer; INL, inner nuclear

layer; NL, merged nuclear layer; GCL, ganglion cell layer. Scale bars, 25 μm. $n = 3$–5 eyes per genotype per age.

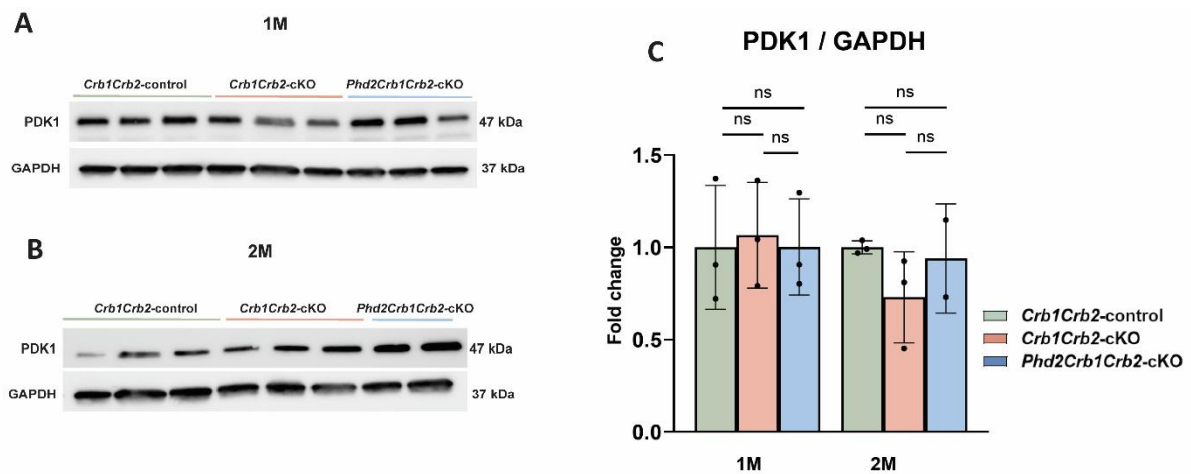

**Supplementary Figure S2. PDK1 protein abundance is not altered following retinal *Phd2* deletion.** Representative Western blots of retinal lysates from *Crb1Crb2*-control, *Crb1Crb2*-cKO, and *Phd2Crb1Crb2*-cKO mice at (A) 1 month and (B) 2 months of age. Retinal lysates were immunoblotted for PDK1 and GAPDH, with GAPDH serving as the loading control. (C) Densitometric quantification of PDK1 protein abundance normalized to GAPDH and expressed as fold change relative to the age-matched control group. No significant differences in PDK1 abundance were detected among the genotypes at either age. Data are presented as mean ± SD; individual points represent biological replicates. At 1 month, $n = 3$ mice per genotype; at 2 months, $n = 3$ for the control and *Crb1Crb2* cKO groups and $n = 2$ for the *Phd2Crb1Crb2* cKO group. Statistical significance was assessed using one-way ANOVA followed by Tukey's multiple-comparisons test. ns, not significant.

**Supplementary Table S1.** Primary antibodies used for this study.

| Antibody | Dilution | Supplier | Identifier |
| --- | --- | --- | --- |
| SOX9 | 1:250 | Millipore | AB5535 |
| Red/Green Opsin | 1:300 | Merck millipore | AB5405 |
| PSD –95 | 1:200 | Cell signaling | 3450S |
| Rhodopsin | 1:100 | Sigma | SAB4502636 |
| PALS1 | 1:100 | Proteintech | 17710-1-AP |
| GFAP | 1:200 | Dako | Z0334 |
| Recoverin | 1:750 | Millipore | AB5585 |
| IBA-1 | 1:250 | Proteintech | 10904-1-AP |
| CD31 | 1:200 | eBioscience | 14-0311-82 |
| GLUT-1 | 1:1000 | Cell Signaling Technology | 12939 |
| LDHA | 1:1000 | Cell Signaling Technology | 2012S |
| PDK1 | 1:1000 | Abcam | EPR19573 |
| GAPDH | 1:2000 | Proteintech | 60004-1-Ig |

**Supplementary Table S2.** Primer sequences for transcriptomic qPCR analysis, related to Figure 8.

| Name | Sequence |
| --- | --- |
| Hif1a – F | CCACAGGACAGTACAGGATG |
| Hif1a – R | TCAAGTCGTGCTGAATAATACC |
| Glut1 – F | GTATGTGGAGCAACTGTGCG |
| Glut1 – R | TCATCGAAGGTTCGGCCTTT |
| Ldha – F | GTCCAGCGAAACGTGAACAT |
| Ldha – R | TCCAAGCCACGTAGGTCAAG |
| Phd2 – F | AGTGACGACGTGTTGCTTCT |
| Phd2 – R | AGACGGGCCAAGAAAAACCT |
| Pdk1 – F | AGGATCGGTTATGCCAACTGT |
| Pdk1 – R | CCAGGAACTTCACACCCCAT |
| Vegfa – F | CCCACGTCAGAGAGCAACAT |
| Vegfa – R | TGCGCTTTCGTTTTTGACCC |
| Hprt – F | CCTAAGATGAGCGCAAGTTGAA |
| Hprt – R | CCACAGGACTAGAACACCTGCTAA |
| Ef1a – F | CTGGATGCTCGCCATCAAA |
| Ef1a -R | GGCGCTTTTCCTCTTGAAGAA |
